# TOR coordinates with transcriptional and chromatin machinery to regulate thermotolerance and thermomemory

**DOI:** 10.1101/2020.07.28.224634

**Authors:** Mohan Sharma, Muhammed Jamsheer K, Brihaspati Narayan Shukla, Manvi Sharma, Prakhar Awasthi, Sanjeet Kumar Mahtha, Gitanjali Yadav, Ashverya Laxmi

**Affiliations:** National Institute of Plant Genome Research, Aruna Asaf Ali Road, New Delhi-110067, India; Amity Food & Agriculture Foundation, Amity University, Uttar Pradesh, India

## Abstract

Global warming exhibits profound effects on plant fitness and productivity. To withstand stress, plants sacrifice their growth and activate protective stress responses for ensuring survival. However, the switch between growth and stress is largely elusive. In the past decade, emerging role of Target of Rapamycin (TOR) has been studied linking energy and stress signaling. Here, we have identified an important role of Glc-TOR signaling in plant adaptation to heat stress (HS). Glc-TOR via the E2Fa signaling module regulates the transcription of heat shock factor genes through direct recruitment of E2Fa onto their promoter regions. Glc also epigenetically governs the transcription of core HS signaling genes in a TOR-dependent manner. TOR acts in concert with p300/CREB HISTONE ACETYLTRANSFERASE1 (HAC1) and dictates the epigenetic landscape of HS loci to regulate thermotolerance. Arabidopsis plants defective in *TOR* and *HAC1* exhibited reduced thermotolerance with a decrease in expression of core HS signaling genes. In addition, TOR also promotes accumulation of histone H3K4me3 marks at the promoters of thermomemory-related genes and therefore, governs thermomemory. Collectively, our findings thus reveal a mechanistic framework in which Glc-TOR signaling through different modules determines the integration of stress and energy signaling to regulate thermotolerance and thermomemory.

## INTRODUCTION

Plants are immobile in nature and are unable to escape the harmful conditions in the environment. Rather, over the course of evolution, they have evolved adaptive strategies to gauge and respond to these stressful conditions for survival and reproductive success. At the time of stress, plants sacrifice their growth to activate protective stress machinery and ensure survival. Once stress signal has vanished, plant quickly cease the stress machinery and activate plant growth and recovery. Sugars produced through photosynthesis are pivotal for plant growth and development. Global transcriptome profiling in response to sugars or mutants defective in sugar signaling (such as TOR) showed alteration in stress induced gene transcripts such as heat shock proteins (Mishra, Singh, Aggrawal & Laxmi 2009; Xiong *et al.* 2013; Gupta, Singh & Laxmi 2015a). TOR is an evolutionary conserved ser/thr kinase, comprised of a multiprotein complex and is a central regulator of different environmental responses in eukaryotes. TOR kinase was first identified in mammals as a target of a molecule named rapamycin isolated from a bacteria *Streptomyces hygroscopicus* (Sehgal, Baker & Vézina 1975). The complex domain architecture of Arabidopsis TOR is evolutionary conserved among eukaryotes including mammals and plants (Menand *et al.* 2002; Zoncu, Efeyan & Sabatini 2011). TOR transduces photosynthesis derived Glc/energy signals to activate progenitor stem cells in the root and shoot apical meristem (RAM and SAM). TOR through interaction and direct phosphorylation, activates E2Fa/b transcription factors in controlling RAM and SAM (Xiong *et al.* 2013; Li *et al.* 2017). E2F-DP controls the transcriptional regulation of genes underlying DNA replication and cell cycle through recognition of conserved cis-acting TTTCCCGCC and other similar elements in their promoter region (Vandepoele *et al.* 2005; Xiong *et al.* 2013). E2Fs activity is controlled by an upstream regulator, Retinoblastoma-related (RBR) tumour suppressor protein. E2Fa in coordination with RBR regulates the balance between proliferation and differentiation (Leviczky *et al.* 2019), however, their roles in stress responses are poorly understood except in replication stress. Interestingly, RBR was also found to be phosphorylated by TOR kinase (Van Leene *et al.* 2019).

In past years, TOR has emerged as a central regulator of nutrient and stress responses. TOR overexpressing (OE) plants showed longer primary roots under excessive nitrate. Moreover, plants with reduced TOR levels showed shorter primary roots with an increased sensitivity to osmotic stress (Deprost *et al.* 2007). Cold exposure caused the transient inhibition of TOR kinase activity and converted the purple hypocotyl of *tor RNAi* lines into green which may be due to decreased translation in *tor RNAi* lines (Wang *et al.* 2016). Under non-stress condition, TOR kinase phosphorylates ABA receptor PYL which dissociates the link between ABA and PP2C phosphatase, conferring inactivation of SnRK2. Onset of stress leads to phosphorylation of RAPTOR1 through ABA activated SnRK2 which disrupts TOR complex association and therefore, inhibition of TOR kinase activity. Plants employ this conserved phospho-regulatory loop to adjust the homeostasis of growth and stress response (Wang *et al.* 2018). In addition to abiotic stresses, TOR negatively regulates defence response to bacterial and fungal pathogens through an antagonistic interaction with jasmonic and salicylic acid signaling (De Vleesschauwer *et al.* 2017).

Plants possess an inherent ability to withstand certain levels of stress, an ability called basal tolerance. In nature, plants are not only challenged by a single exposure of stress, but stresses are chronic and sometimes irregular. Thus, plants have evolved strategies that enables them to withstand stress in a superior manner if prior confrontation with the same stress, called acquired tolerance. The preservation of acquired tolerance in the leisure of interfering stress is called stress memory (Balmer, Pastor, Gamir, Flors & Mauch-Mani 2015). Numerous reports have described stress memory in response to salt, drought, heat and pathogen (Conrath 2011; Ding, Fromm & Avramova 2012a; Sani, Herzyk, Perrella, Colot & Amtmann 2013; Lämke, Brzezinka, Altmann & Bäurle 2016; Sharma, Banday, Shukla & Laxmi 2019). The majority of stress responses revert back to pre-stress stage shortly after stress, which allows them to forget stressful period and allocate resources to support plant growth. In contrast, stress memory enables plants to respond more efficiently to future stress suggesting essential temporal dynamics but underexplored.

Chromatin structures lie at the core of stress responses and play crucial roles ranging from sensory to downstream gene activation (Kumar & Wigge 2010; Lämke & Bäurle 2017). Structure of chromatin is remodelled at various levels such as alternation in histone variants, nucleosome positioning, gene looping and post-transcriptional histone and protein modifications (Deal & Henikoff 2010; Luo *et al.* 2012; Struhl & Segal 2013). In most cases, modification of histone lysine through acetylation correlates with active gene transcription whereas lysine methylation at different positions serve as activating or repressing marks, for example, H3K4me2 and H3K4me3 promote active gene transcription whereas H3K27me3 serve as repressive histone mark. In addition, histone demethylation serves as important determinant of chromatin structures and performs critical roles in developmental and stress responses (Cui *et al.* 2016; Liu *et al.* 2019). The member of SWI/SNF complex BRAHMA (BRM) acts in concert with H3K27me3 demethylase REGULATOR OF EARLY FLOWERING 6 (REF6) and performs active roles in gene transcription at the chromatin level. REF6 facilitates the recruitment of BRM at specific genomic loci containing a CTCTGYTY motif (Li *et al.* 2016). Recent report demonstrated an important role of BRM and REF6 in thermomemory and transgenerational thermomemory (Brzezinka *et al.* 2016; Liu *et al.* 2019). BRM defective plants displayed reduced sustenance of thermomemory-associated gene expression, leading to reduced thermomemory (Brzezinka *et al.* 2016).

Lately, Sharma et al., 2019 identified the pivotal role of Glc and Glc-activated TOR signaling in plant response to HS. Arabidopsis TOR overexpressing plants showed increased short term acquired thermotolerance (SAT) whereas plants with subdued TOR displayed reduced SAT (Sharma *et al.* 2019). However, the underlying mechanism how TOR controls thermotolerance response is unknown. Here, we show important regulatory role played by Glc-TOR in mediating thermotolerance response. Whole genome transcriptome analysis of Arabidopsis seedlings in response to heat and Glc sufficiency exhibited large synergistic interaction with TOR transcriptomic data. TOR, in concert with p300 histone acetyltransferase 1 (HAC1) modulates the acetylation of HS loci in conferring SAT. In addition to SAT, Arabidopsis TOR remembers the past exposure of HS for several days through maintenance of expression and protein levels of thermomemory associated gene(s). TOR promotes the accumulation of H3K4me3 methylation at the promoters of thermomemory-related genes and thus causing thermomemory.

## MATERIAL AND METHODS

### Plant Materials and Growth Condition

The *Arabidopsis thaliana* mutant lines estradiol-inducible TOR RNAi (CS69829; *tor-es1*; Col-0 background), TOR OE (AT1G50030, GK-548G07-020632; Col-0 background) and *e2fa* (AT2G36010, WiscDsLox434F1; Col-0 background) were obtained from ABRC at Ohio State University. The *TOR* (AT1G50030, G166; Col-0 background) line was obtained from the NASC. The following lines were obtained from the published sources: *tor 35-7* RNAi (Col-0 background (Deprost *et al.* 2007) and *HAC1* mutants (*hac1-2*, *hac1-3* and *hac1-6*; Col-0 background (Heisel, Li, Grey & Gibson 2013), *proTOR::GUS* (Col-0 background (Menand *et al.* 2002). Seeds were sown on square petri plates (120*120mm) containing 0.5% MS medium with 1% Suc (29mM) and 0.8% agar (24mM), under long-day photoperiod (16h light and 8h dark, 60mM m^−2^ s^−1^ light intensity) at 22°C±2°C temperature.

### GUS Assay and Chl estimation

GUS assay was performed using seven-day-old *proTOR::GUS* reporter transgenic line treated without or with Glc (−/+ Glc) followed by HS. HS was applied as 1 h at 37°C, 2 h at 22°C, and 2.5 h at 45°C. After HS, *proTOR::GUS* seedlings were transferred immediately (0h) after 24h of HS recovery at 22°C to GUS buffer. Images were taken using stereo zoom microscope (GUS treatment was described by Mishra et al., 2009). Total Chl (Chl a and Chl b) estimation was performed with seedlings treated with HS. Chl estimation was described by (Kushwah & Laxmi 2014).

### ChIP Assays

ChIP assays were performed using protocol described by Saleh et al. (2008) with minor modifications. Protoplasts were harvested from Col-0 plants and treated with Glc (5mM) for 3 h followed by chromatin isolation. For ChIP assay in seedlings, seven-day-old Arabidopsis seedlings were starved with Glc free medium for 24 h followed by 3h of Glc treatment. For ChIP assay used in thermomemory, seven-day-old Arabidopsis seedlings were transferred directly to Glc-containing media followed by HS treatment. HS was applied as 1 h at 37°C, 1.5 h at 22°C, and 45 min at 45°C. Following HS, seedlings were recovered for 24h at 22°C. The resuspended chromatin was sonicated in a 4°C water sonicator (Diagenode Bioruptor Plus). Serum containing anti-E2Fa antibodies was obtained from Prof. Lieven De Veylder, VIB Department of Plant Systems Biology, Ghent University (Takahashi *et al.* 2008). Antibodies against H3K (9+14+18+23+27) acetylation were purchased from Abcam (Cat. no. ab47915). Antibodies against H3K4me2 (Cat. no. 07-030) and H3K4me3 (Cat. no. 07-473) were purchased from Millipore. All primers used are mentioned in the Supplemental Table S7.

### Western blot Assay

The total plant protein from seven-day-old Arabidopsis seedlings was extracted in cold extraction buffer (137mM of NaCl, 2.7mM of KCl, 4.3mM of Na2HPO4, 1.47mM of KH2PO4, 10% glycerol [1.1 M], and 1mM phenyl methyl sulfonyl fluoride [PMSF]) together with plant protease inhibitor cocktail (Sigma-Aldrich, http://www.sigmaaldrich.com/). Protein concentration was measured using nanodrop. 50 µg of protein samples were loaded in each well. The antibody for HSP21 (Anti-HSP21; chloroplastic heat shock protein Cat. no. AS08 285) and (HSP90-2; Cat. no. AS11 1629) were purchased from Agrisera.

### Thermotolerance/thermomemory assays

*Arabidopsis thaliana* seedlings grown in standard culture room and maintained under long day photoperiod conditions were used for thermotolerance assays. Seven-day-old seedlings were grown initially in MS media using 1% sucrose (29mM) and 0.8% Agar and then accimatized to Glc free and Glc (167mM) containing solid MS media for 24 h followed by SAT and LAT HS treatments. For SAT and LAT treatment, seedlings were kept in HS as 37°C for 1h, allowed to recover for 2h (SAT) and 2d (LAT) at 22°C and then transferred to 45°C for 2.5h, followed by recovery in the culture room for 3–4d. To estimate the length of thermomemory, seven-day-old seedlings were transferred to the heat chamber at 37°C for 3h, allowed to recover for 7d at 22°C and then transferred to 45°C for 3h, followed by recovery in the culture room for 3–4d. For immunoblotting assay, seven-day-old Arabidopsis seedlings were transferred to HS at 37°C for 1h, 22°C for 90min, 45°C for 45min followed by recovery 22°C for 2d, 5d and 7d.

### Gene expression analysis

For RT-qPCR study in response to Glc, 5-d-old Arabidopsis seedlings were starved without Glc containing MS medium for 24h in dark followed by without or with Glc containing MS medium for 3h in light. For RT-qPCR study under thermomemory HS, Arabidopsis seedlings were germinated and grown directly in medium containing 1% Glc (56mM) for seven days under standard 16-h day/8-h day night conditions. For SAT and LAT, seedlings were transferred to HS at 37°C for 1h, 22°C for 2h (SAT) and 2d (LAT), 45°C for 45 min and 22°C for 3h, 24h and 48h. Total plant RNA was isolated using RNAeasy Plant Mini Kit (Qiagen) and cDNA was prepared using 2 µg of total RNA with a high-capacity cDNA Reverse Transcription Kit (Applied Biosystems). All candidate gene primers were designed using a unique sequence within the genes preferably 3’UTR. Normalization of genes was performed using 18S as a reference control. The fold-change for each candidate gene in different experimental conditions was determined using the quantitative DDCT method. All primers used are listed in the Supplemental Table S7.

### Microarray Analysis

Whole genome microarray analysis of five-day-old Arabidopsis Col-0 seedlings treated without or with Glc (−/+ Glc) and HS was performed. To minimize the endogenous sugar background, five-day-old Col-0 seedlings were first starved with ½ MS medium without sugar for 24h in the dark followed by 24h Glc acclimation and then subjected to HS. HS treatment was applied as 37°C for 3h and 37°C for 3h followed by recovery at 22°C for 3h. Harvested samples were outsourced for transcriptome profiling. The data was analysed using the Transcriptome Analysis Console (v3.0; Affymetrix) using default parameters. DEGs in all other conditions were compared to -Glc with fold change (FC) value (>= to 1.5). One-Way Between-Subject ANOVA (unpaired) statistical test with p-value less than 0.05 and false discovery rate (FDR) p-value 0.1 were used to analyse the DEGs. Microarray data has been submitted to Array Express with accession no. E-MTAB-6980. Three biological replicates were used in this microarray study.

### Bioinformatics analysis

BRM/REF6 promoter motifs in TOR co-expression genes, *tor_DEG* and E2Fa motif containing genes were extracted for a length of 1200 nucleotides (including an upstream region of 1000 nts and downstream region of 200 nts, from the TSS sites). These regions were then scanned for the presence of eight nucleotide length motif “CTCTGYTY” (where Y is any one of A/T/G/C) and this was done using in-house python scripts. Gene expression data for TOR, RAPTOR1, HAC1 and BRM in *A. thaliana* across various anatomical parts, developmental stages and stress related experiments was extracted using the mRNA-Seq dataset ‘AT_mRNASeq_ARABI_GL’ from Genevestigator platform (Hruz *et al.* 2008). The available time series expression data (in Transcripts per million TPM counts) for Arabidopsis diurnal range across the day-night cycle was downloaded from the Supplementary dataset in Ferrari et al., 2019 (Ferrari *et al.* 2019). Temporal expression patterns for all four genes were extracted from this dataset and compared using Normalized TPM values (rescaled between 0 and 1 via Min-Max scaling/normalization). Expression Heat maps were generated using MeV_4_9_0.

### Accession Numbers

Sequence data from this article can be found in the GenBank/EMBL data libraries under accession numbers: AT1G50030 (TOR), AT1G79000 (HAC1), AT2G36010 (E2FA), AT4G17750 (HSFA1A), AT2G26150 (HSFA2), AT3G12580 (HSP70), AT1G54050 (HSP20-LIKE), AT1G52560 (HSP20-LIKE), AT4G27670 (HSP21), AT5G59720 (HSP18.2), AT5G12020 (HSP17.6II), AT4G10250 (HSP22), AT1G74310 (HSP101) and AT3G18780 (ACT2).

## RESULTS

### Exogenous glucose mimics the natural environmental conditions in governing thermotolerance

Glc is a vital source of energy for most of the organisms on earth. It serves as a fundamental metabolic and signaling molecule in regulating multifaceted roles including transcription, translation, cell cycle, disease and stress responses. Recently, Sharma et al., 2019 revealed the important role of Glc signaling in thermotolerance. In addition, Glc activated TOR-E2Fa signaling is required to cope deleterious effects of HS (Sharma *et al.* 2019). However, the molecular mechanism how Glc driven TOR signaling governs temperature mediated response is hitherto unknown. Under natural environment, glucose production depends on the availability of light. This photosynthetically generated high glucose levels could modulate root architecture and endogenous sugar levels are regulated by fluctuating light conditions (Kircher & Schopfer 2012; Gupta, Singh & Laxmi 2015b). To test whether high light (HL) generated glucose could provide thermotolerance to Arabidopsis plants, we grew plants under low (LL; 2000 lux) and high light (10000 lux) in presence of low and moderate Glc and effect of HS was analysed. HS was applied as 1h_37ºC, 2h_22ºC, 2.5h_45ºC and 3-7d_22ºC. HL intensity (10000 lux) could increase the HS tolerance in Arabidopsis seedling as compared to LL (2000 lux) (Figure 1A-D). HL along with moderate glucose (56mM) displayed synergistic effect on thermotolerance, as shown by robust plant growth (Figure 1A-D). Earlier, Sharma et al., 2019 also showed that plants growing under Glc (3%Glc; 167mM) showed enhanced survival at HS. We propose that 1%Glc with HL is similar to previously reported conditions and could mimic optimum Glc response (3%Glc) for plants to survive under HS (Sharma *et al.* 2019). Next, we assessed the phenotype of Arabidopsis Col-0, TOR OE lines (G166 and G548) and *tor35-7* RNAi lines under HL and LL in presence of low and moderate Glc and challenged them for HS. Arabidopsis TOR OE survived better under HL than Col-0 plants. In contrast, *tor35-7* RNAi exhibited severe death phenotype under HL (Figure 1A-D). In addition, TOR OE displayed higher growth recovery as revealed by higher fresh weight, chlorophyll content and increased number of lateral roots under HL whereas *tor35-7* showed reduced growth and recovery (Figure 1A-C). Since photosynthesis produces sucrose that is converted to glucose and fructose before entering metabolism, it was important to test whether HL-mediated thermo-protection is specific to Glc. To test this, we blocked Glc signaling/metabolism by using Glc analogue 2-deoxy-Glucose (2-DG) and checked thermotolerance response under HL. 2-DG is actively transported through hexose transporters and phosphorylated but cannot be metabolised. 2-DG-phoshate interferes with Glc metabolism through inhibition of glycolytic enzymes (Ralser *et al.* 2008). Five-day-old MS grown Arabidopsis Col-0 seedlings under normal light intensity were transferred to Glc (56mM) containing MS media without or with 2-DG (2.5mM). After transfer, seedlings were transferred to HL for 24h and then subjected to HS. Interestingly, we found that plants treated with 2-DG could not provide tolerance under HL further substantiating specific role of photosynthesis derived Glc signaling in governing thermotolerance (Supplemental Figure 1A-D).

**Figure 1.**
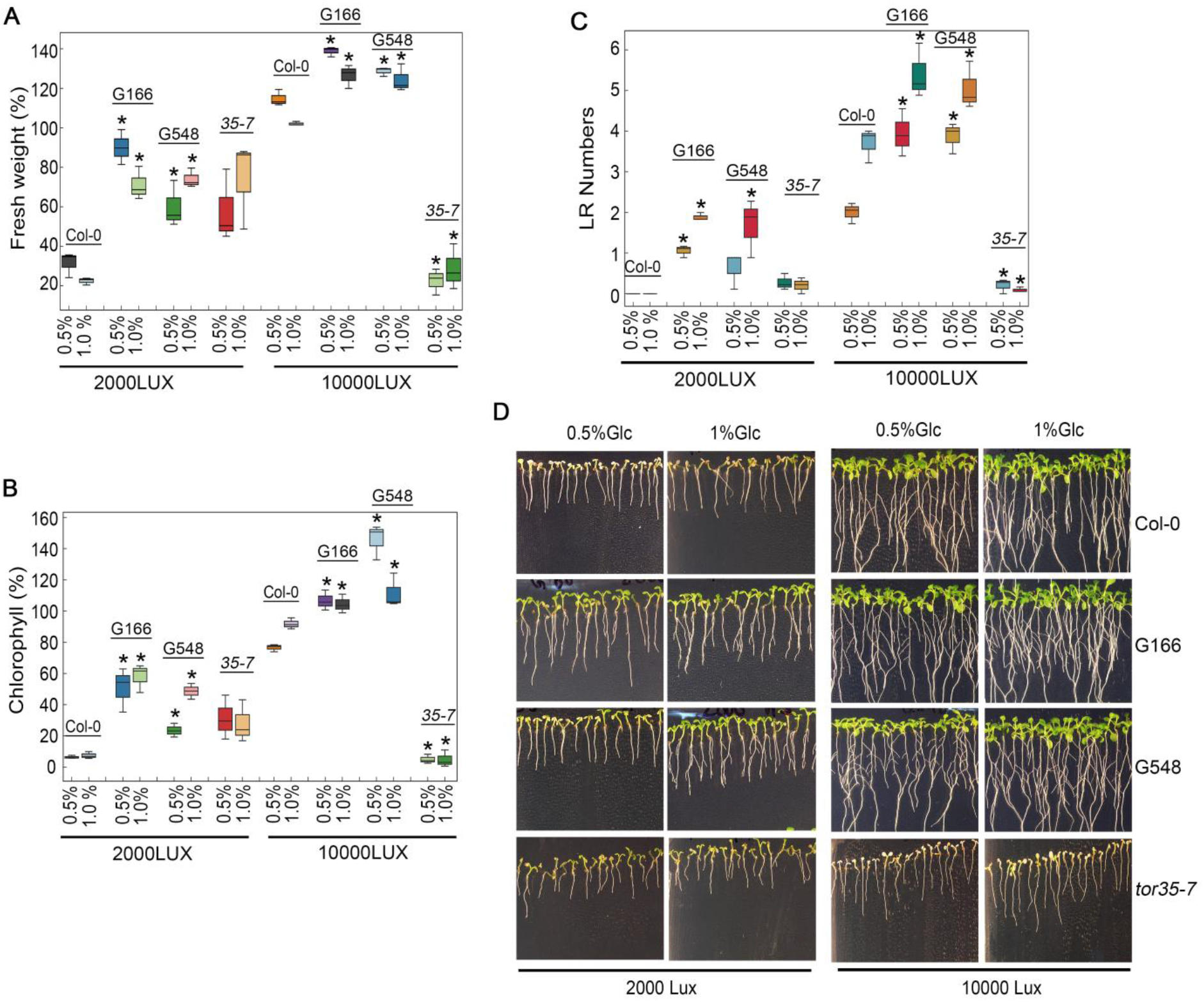
Increasing light intensity mimics Glc-TOR mediated thermotolerance response. **A-C**, Percentage measurement of fresh weight, Chl content and LR numbers of Arabidopsis Col-0 TOR overexpression (G166 and G548) and *tor 35-7* RNAi lines grown under low (2000LUX) and high (10000LUX) light intensity after HS. **D**, HS phenotype of Arabidopsis Col-0, G166, G548 and *tor 35-7* lines grown under low (2000LUX) and high (10000LUX) light intensity. Arabidopsis seedlings were grown directly onto MS medium containing 0.5% (28mM) and 1% (56mM) Glc for seven days under low and high light intensity and then transferred to HS. HS was applied as 1h_37ºC, 2h_22ºC, 2.5h_45ºC followed by recovery at 3-4d_22ºC under same low (2000LUX) and high (10000LUX) light intensities. Data shown are average of three biological replicates. Experiments were repeated thrice with similar results. Error bars represent SD. (Student’s t test, P < 0.05; *WT versus mutant/overexpression).

### Presence of Glc profoundly alters the expression of HS regulated genes

To dissect the contribution of Glc/energy in HS transcriptional response, whole genome microarray analysis of Arabidopsis Col-0 seedlings was performed. Five-day-old seedlings were subjected to Glc free (-Glc) MS media for 24h followed by 24h Glc treatment (+Glc; 167mM). Following acclimation to Glc medium, seedlings were transferred to HS (37°C_3h) and HS Rec (37°C_3h followed by 22°C_3h). The cRNA was prepared from RNA isolated from Glc and HS treated samples and were hybridized onto Affymetrix Arabidopsis whole genome ATH1 arrays. The extent and overlap of Glc and HS regulated genes at the transcriptional level was analysed to determine the unique and overlapping set of genes, which are crucial targets of Glc-HS signaling crosstalk. We used six treatment conditions and designated following terms for Glc free (-Glc), Glc only (+Glc), HS only (-Glc/+HS), Glc and HS (+Glc/+HS), HS followed by recovery (-Glc/+HS Rec) and Glc and HS followed by recovery (+Glc/+HS Rec).We considered the differential expression of genes (DEG) affected 1.5-fold or more by HS and Glc. Out of 2311 HS (-Glc/+HS) DEGs, 1124 (49%) genes were also found to be up- and down-regulated by Glc and HS (+Glc/+HS; Figure 2A; Supplemental Table S1A). Among 1124 genes, 663 (59%) genes were synergistically (down-regulated by both −Glc/+HS and +Glc/+HS) downregulated and 452 genes were synergistically up-regulated (up-regulated by both categories) (Figure 2B and 2C; Supplemental Table S1A). Only 9 (0.8%) genes were antagonistically (up in one category and down in other and vice versa) regulated by both –Glc/+HS and +Glc/+HS. The other feature of interest was to analyse how the –Glc/+HS mediated induction or suppression is affected in seedlings treated with +Glc/+HS. Among 1124 –Glc/+HS and +Glc/+HS co-regulated genes, the extent of HS induction/suppression was affected (FC ≥ +/− 1.5) in 259 (24%) genes in the presence of Glc (Figure 2D; Supplemental Table S1B). Out of these 259 genes, Glc was able to affect the extent of HS regulation antagonistically in 174 (115+59; 67%) genes and agonistically in 85 (65+20; 33%) genes. Out of 460 –Glc/+HS up-regulated genes, Glc could affect the HS induction in 180 (115+65; 39%) genes of which 115 (64%) genes were affected antagonistically and only 65 (36%) genes were affected agonistically. Similarly, out of 664 −Glc/+HS down-regulated genes, presence of Glc could affect the extent of HS mediated suppression in 79 (59+20; 19%) genes. Out of these 79 genes, presence of Glc could affect 59 (75%) genes antagonistically while only 20 (25%) genes were agonistically regulated by Glc (Figure 2D). Interestingly, an exclusive set of genes was also regulated by both –Glc/+HS (971 genes) and +Glc/+HS (1408 genes) category (Supplemental Table S1C and S1D). GO up-regulated categories enriched under –Glc/+HS conditions mostly belonged to stress responses whereas growth related processes such as root, embryo, seed and fruit development, fatty acid, lipids and amino acid biosynthetic processes were severely down-regulated (Figure 2E and Supplemental Figure S3A). In contrast, processes related to nucleotide biosynthesis, rRNA processing and ribonucleoprotein complex biogenesis required for ribosome synthesis were highly up-regulated only in +Glc/+HS (Figure 2E). TOR, as a master regulator, senses and transduces cellular energy signals and regulates transcriptome reprogramming of myriad genes (Xiong *et al.* 2013). We, therefore, compared our transcriptome data with publicly available Glc/TOR transcriptome of three day-after-germination (DAG) seedlings of WT and *tor* treated with Glc for 2h (Xiong *et al.* 2013). We found that Glc-TOR and +Glc/+HS regulate transcriptome reprogramming largely by synergistic interaction (total 560 genes; 310 genes agonistically down-regulated and 250 genes agonistically up-regulated by both Glc-TOR and +Glc/+HS) and only a small number of genes were regulated antagonistically (Figure 2F; Supplemental Table S2A). GO categories enriched under synergistic down-regulation were involved in sucrose starvation and catabolic processes whereas protein folding, de-novo posttranslational protein folding, peptidyl-prolyl isomerization and protein complex oligomerization were synergistically up-regulated (Supplemental Figure S3B and S3C). Surprisingly, –Glc/+HS category showed major antagonism with Glc-TOR microarray (total 312 genes; 173+139 genes; Figure 2G; Supplemental Table S2B, Supplemental Figure S4A and S4B). In contrast, 176 (97+79) genes were agonistically regulated by both Glc-TOR and –Glc/+HS (Figure 2G; Supplemental Table S2B, Supplemental Figure S4C and S4D). Collectively, these results suggest that Glc regulates the large transcriptome of HS genes in a TOR-dependent manner.

**Figure 2.**
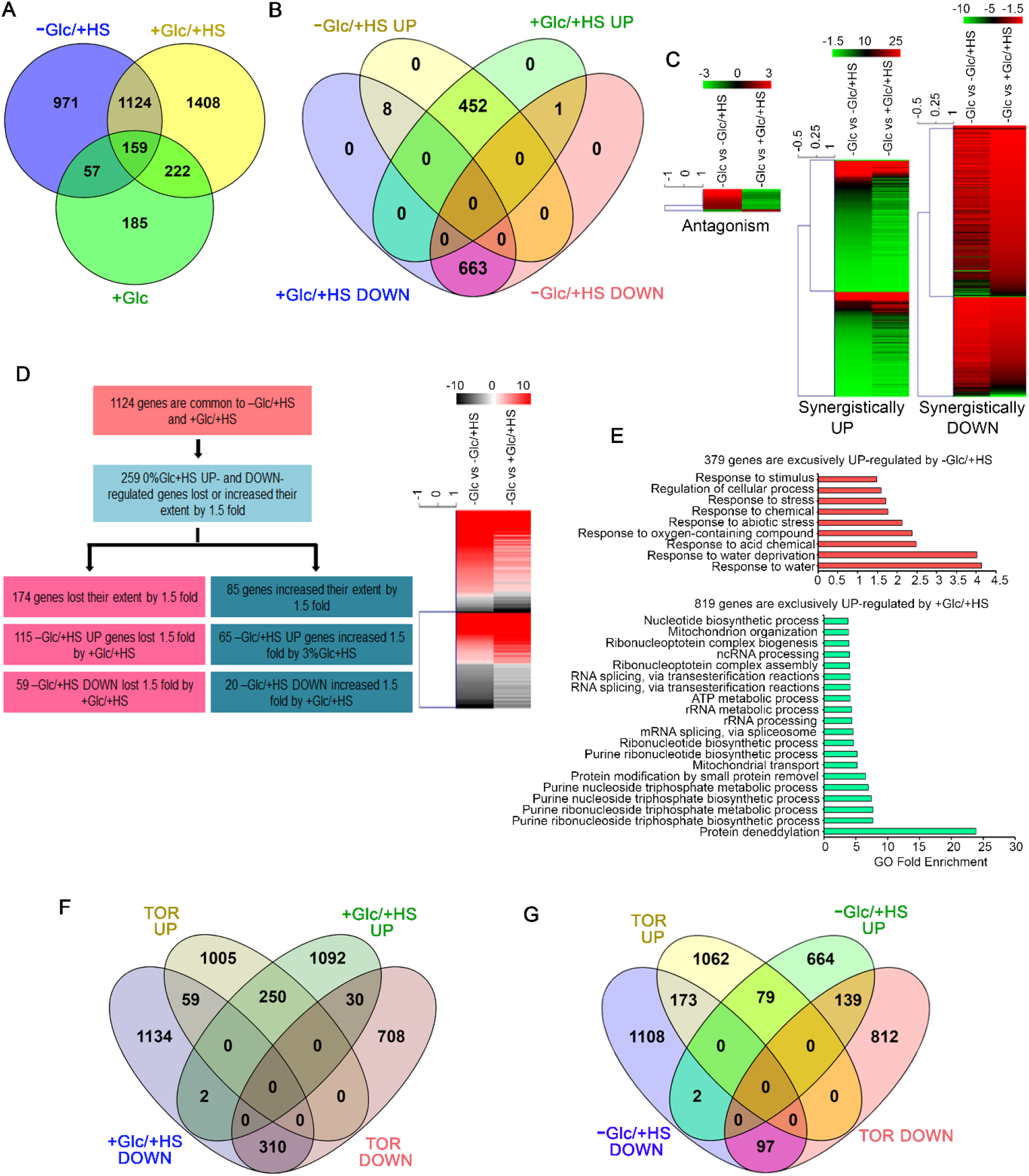
Glc triggers transcriptome reprogramming of HS-regulated genes. **A**, Venn diagram showing overlap of genes induced by –Glc/+HS, +Glc/+HS and +Glc only. **B, C**, Venn diagram and heat maps showing synergistic and antagonistic relation of genes induced by –Glc/+HS and +Glc/+HS. **D**, Picture showing difference in gene expression extent in common gene affected both by –Glc/+HS and +Glc/+HS. Genes induced by –Glc/+HS affected (lost/increase in extent) by 1.5 fold in presence of Glc was taken into consideration. **E**, Gene ontology analysis of biological process enrichment of genes exclusively induced by –Glc/+HS and +Glc/+HS, respectively. Top twenty GO categories were plotted. **F**, Venn diagram showing synergistic and antagonistic interaction of +Glc/+HS transcriptome with publicly available Glc/TOR transcriptome of three-day-old seedlings of WT and *tor* treated with Glc for 2h. **G**, Venn diagram showing synergistic and antagonistic interaction of –Glc/+HS transcriptome with publicly available Glc/TOR transcriptome. Data shown are average of three biological replicates. Venn diagrams were created using venny v2.1, heat maps were created using Mev and GO biological process were created using Panther GO term enrichment tool.

### Glc triggers HS transcriptome for better growth and recovery after harmful stress

To understand how Glc regulates HS response under post stress period, we analysed the overlap of HS transcriptome with post-HS (HS followed by recovery at 22°C) transcriptome data. We compared the genes regulated by +Glc/+HS with that of +Glc/+HS Rec. Intriguingly, we observed that Glc maintained the sustenance of expression in 1332 (702+630) genes (Figure 3A; Supplemental Table S3A). Functional categorization of 702 genes co-regulated by +Glc/+HS and +Glc/+HS Rec enriched many important biological processes required for plant protection and HS recovery including protein folding/refolding, rRNA metabolism and processing, rRNA transcription and ribosome biogenesis whereas processes related to sucrose starvation and catabolism were down-regulated (Figure 3C and 3E, Supplemental Table S3B, Supplemental Figure S5A). Stress response leads to cessation of plant growth and once the stress has gone, plants regain their growth. Growth requires cell division and proliferation from meristematic organs. Remarkably, 630 genes were exclusively regulated by +Glc/+HS Rec with enrichment of processes related to cell division, cell wall modification, DNA unwinding, geometric changes and replication together with histone/protein methylation indicating a major role of Glc/energy in post-stress growth recovery which was abrogated in heat-stressed plants lacking Glc/energy (Figure 3A and 3B; Figure 3F; Supplemental Table S3C). Astonishingly, out of 2311 –Glc/+HS differentially regulated genes, only 19 (0.8%) genes showed their sustained expression when recovered at 22°C without Glc. (Figure 3B and 3D; Supplemental Table S3D and S3E).

**Figure 3.**
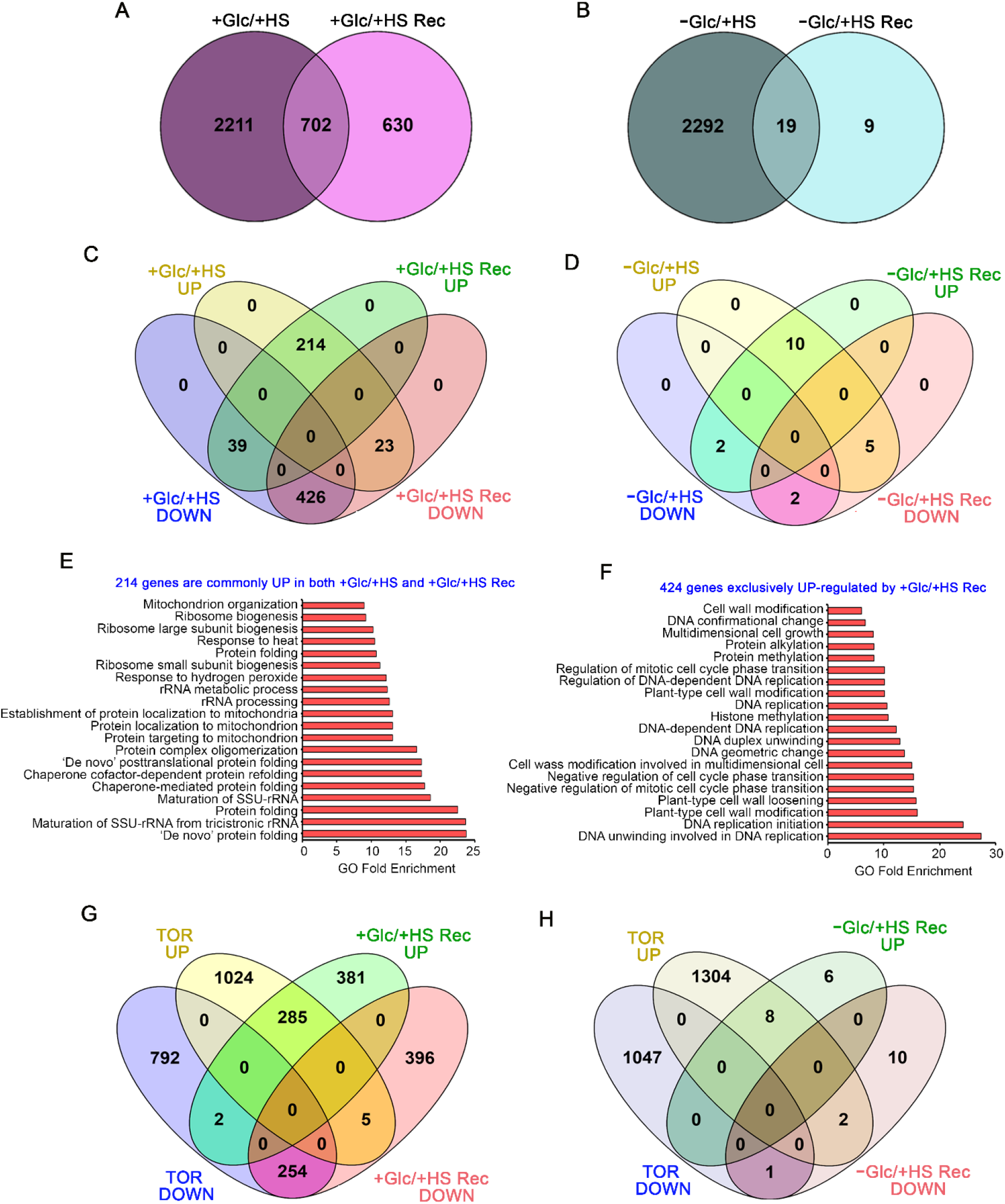
Presence of Glc is obligatory for transcriptome reprogramming of HS recovery. **A**, Venn diagram showing overlap of genes induced by +Glc/+HS and +Glc/+HS Rec. **B**, Venn diagram showing overlap of genes induced by –Glc/+HS and −Glc/+HS Rec. **C**, Venn diagram showing synergistic and antagonistic relation of genes induced by +Glc/+HS and +Glc/+HS Rec. **D**, Venn diagram showing synergistic and antagonistic relation of genes induced by –Glc/+HS and −Glc/+HS Rec. **E**, Gene ontology analysis showing biological process enrichment of genes commonly induced by +Glc/+HS and +Glc/+HS Rec. **F**, Gene ontology analysis showing biological process enrichment of genes exclusively induced by +Glc/+HS and +Glc/+HS Rec. **G, H**, Venn diagram showing synergistic and antagonistic interaction of +Glc/+HS Rec and –Glc/+HS Rec transcriptome with publicly available Glc/TOR transcriptome. Data shown are average of three biological replicates. Venn diagrams were created using venny v2.1 and GO biological process were created using Panther GO term enrichment tool.

Next, we compared the transcriptome of −Glc/+HS Rec and +Glc/+HS Rec with Glc-TOR transcriptome data. We observed a large synergistic interaction between Glc-TOR and +Glc/+HS Rec (total 539 genes; 254 genes were synergistically down and 285 up-regulated by both Glc-TOR and +Glc/+HS Rec), while very few genes (7 genes) were antagonistically regulated (Figure 3G; Supplemental Table S2C). Functional categorisation of 285 up-regulated genes showed emergence of peptide and histone arginine methylation together with processes related to DNA replication, RNA modification, rRNA processing and protein folding (Supplemental Figure S6A). Arginine methylation is a most common posttranslational modification functioning as an activator or a repressor of gene transcription (Liu, Lu, Cui & Cao 2010; Fulton, Brown & Zheng 2018). In addition, processes such as sucrose starvation and catabolism were commonly down-regulated by +Glc/+HS and Glc-TOR (Supplemental Figure S6B). Moreover, we checked whether Glc-TOR transcriptome exhibits any overlap with –Glc/+HS Rec. Only very few genes were found to be overlapping with –Glc/+HS Rec and Glc-TOR, further suggesting prerequisite requirement of Glc mediated signaling to cope with lethal HS (Figure 3H; Supplemental Table S2D). Taken together, these results suggest that Glc-TOR and +Glc/+HS Rec signaling governs synergistic regulation of HS transcriptome and therefore, Glc-TOR signaling pathway is obligatory for plants survival under extreme HS and post-stress period for improved growth and recovery.

### TOR target E2Fa binds directly to HSFA1 gene promoters to regulate thermotolerance

In our previous report, TOR target E2Fa binds directly to *HLP1* promoter to activate HS gene expression (Sharma *et al.* 2019). Moreover, genes which are shown to play roles in stress and energy signaling demonstrated E2Fa binding sites in their promoters (Ramirez-Parra, Fründt & Gutierrez 2003; Vandepoele *et al.* 2005; Xiong *et al.* 2013). E2Fa is a transcription factor and is a direct downstream target of TOR which undergoes phosphorylation through TOR upon sugar/energy sufficiency (Xiong *et al.* 2013). It has previously been shown that E2Fa motifs are present in gene promoters of HSFA1 class of HSFs (Ramirez-Parra *et al.* 2003; Naouar *et al.* 2009; Xiong *et al.* 2013). To investigate whether E2Fa directly regulates heat shock gene transcription by binding to their promoters, we performed *in vivo* ChIP using protoplasts from four-week old Col-0 plants. Anti-E2Fa serum was used to immunoprecipitate E2Fa/promoter and approximately 200-250 bp *HSFA1s* promoter comprising E2Fa binding sites were analysed using qPCR. Two or more regions containing similar or identical E2Fa motifs were chosen. The Amount of chromatin enriched by anti-E2Fa was significantly higher in all four *HSFA1s* (*HFSA1a*, *HSFA1b*, *HSFA1d* and *HSFA1e*; Figure 4A-D). Interestingly, E2Fa also binds directly to *HSFA2* gene promoter (Figure 4E). Furthermore, transcript levels of all four *HSFA1 and HSFA2* genes were significantly reduced in *e2fa-1* [Figure 4F; (Sharma *et al.* 2019)]. These findings thus indicate that E2Fa transcriptionally activates *HSFA1s* and *HSFA2* genes by binding directly to their promoters, suggesting crosstalk between TOR-E2Fa and HSFA1/HSFA2 mediated HS signaling.

**Figure 4.**
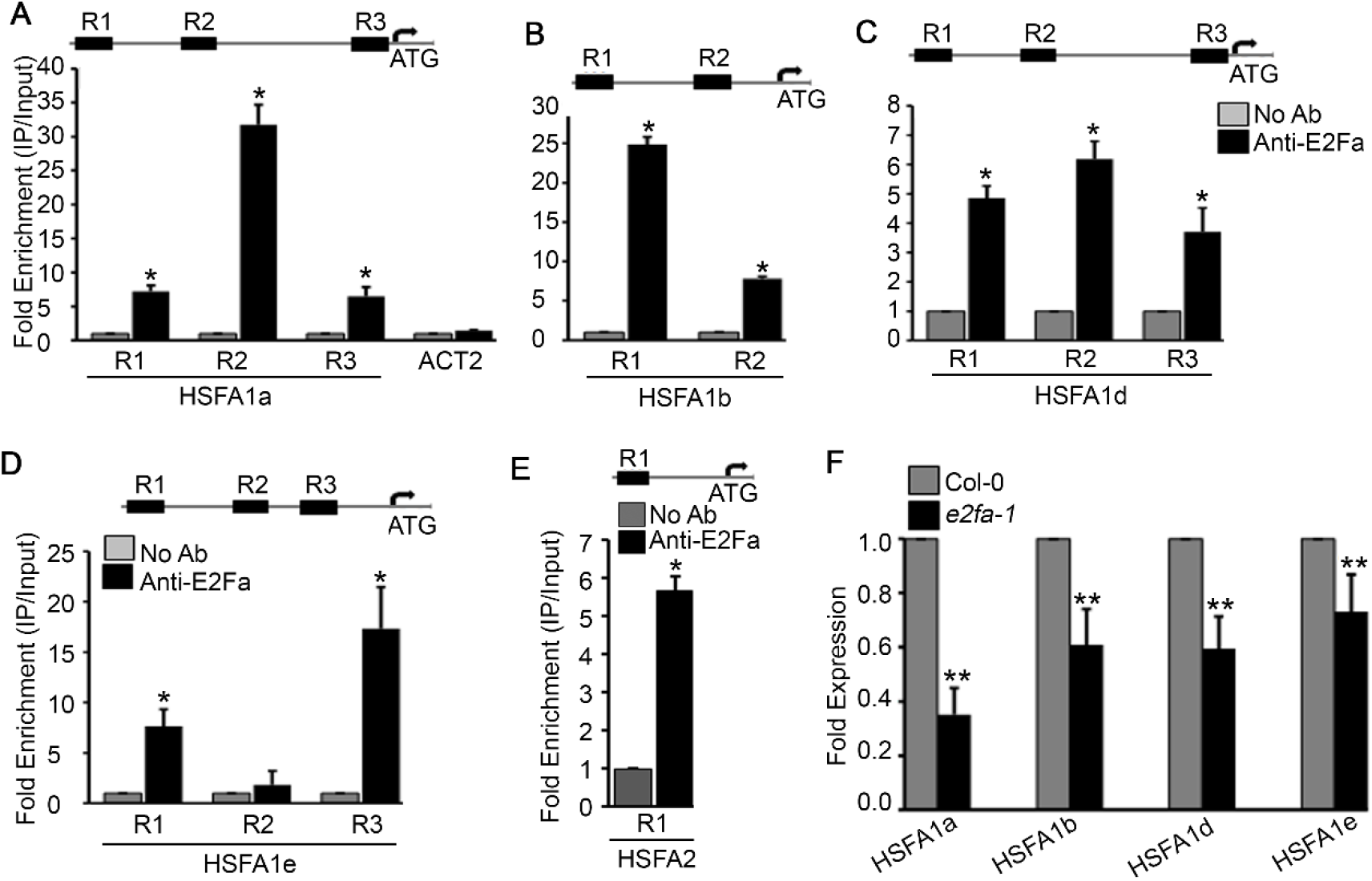
TOR phospho-substrate E2Fa binds directly to the promoters of HSF genes. **A**, ChIP assay of HSFA1a promoter. Amplicon position relative to ATG are as follows: R1 (−1480 to −1281); R2 (−913 to −712) and R3 (−183 to −2). ACT2 was used as a negative control in the ChIP assay. **B**, ChIP assay of HSFA1b gene promoter. Amplicon position relative to ATG of HSFA1b: R1, −1103 to −858; R2, −556 to −351; **C**, ChIP assay of HSFA1d promoter. Amplicon position relative to ATG of HSFA1d: R1, −1381 to −1176; R2, −1031 to −832; R3, −256 to −57. **D**, ChIP assay of HSFA1e promoter. Amplicon position relative to ATG of HSFA1e: R1, −1397 to −1185; R2, −894 to −705; R3, −606 to −404. **E**, ChIP assay of HSFA2 promoter. Amplicon position relative to ATG of HSFA2: R1, −823 to −614. **A-E**, Protoplasts were harvested from four-week-old Col-0 plants. Fold enrichment of promoter fragments were calculated by comparing samples treated without or with Anti-E2Fa specific serum. Schematic representation of 1.5 kb upstream region of HSFA1 and HSFA2 promoter containing E2Fa binding motifs. CT values without and with anti-E2Fa serum were normalized by input control. Data shown are average of four technical replicates among two biological replicates. Experiments were repeated four times with similar results. **F**, RT-qPCR-PCR showing expression of *HSFA1* genes in *e2fa-1* mutant. Five-day-old Arabidopsis Col-0 and *e2fa-1* seedlings were used. Data shown are representative of one biological replicate. Experiments were repeated twice with similar results. Error bars represent SD. (Student’s t test, P < 0.05; *, No Ab vs Ab; **, WT vs mutants).

### Glucose epigenetically regulates the transcription of HS genes in Arabidopsis

According to previous microarray reports, glucose controls the transcription of a large number of heat shock proteins (HSP) genes (Mishra *et al.* 2009; Xiong *et al.* 2013; Gupta *et al.* 2015a). To confirm these microarray results, we performed quantitative real time PCR (RT-qPCR) using nine-day-old MS grown Arabidopsis Col-0 seedlings, starved without glucose for 24h and then treated with glucose (167mM) for 3h. We assessed the expression of five key HS genes. Glucose highly induced the expression of these core HS genes than plants without glucose (Figure 5A). Previously, Sharma et al., 2019 showed that plants growing under various sugars and sugar analogues (3%Glc, 3%mannitol and 3% 3-O-methyl-d-glucopyranose) exhibited specific role of Glc in thermotolerance. They also showed that Glc and Suc specifically induced the expression of heat-related protein genes which further ruled out the possibility of cross-protection due to increased Glc concentration (Sharma *et al.* 2019). Next, we examined HSP expression in Arabidopsis TOR estradiol-inducible RNAi line (*tor-es1*) under glucose sufficiency. To this end, five-day-old MS grown Arabidopsis *tor-es1* seedlings were transferred to estradiol (20µM) containing MS media and kept for four days. Following estradiol treatment, seedlings were starved without glucose for 24h and then treated with glucose (167mM) for 3h. Arabidopsis *tor-es1* showed more than 50% reduction in HS genes expression under glucose+estradiol treatment than mock (DMSO) (Figure 5A). These results suggest that glucose regulates HS gene transcription in a TOR-dependent manner. As Glc regulates the transcripts of the HS genes through the TOR, it was interesting to investigate how Glc regulates this response. To this end, we assessed the promoters of heat shock genes. Histone and chromatin modifications are important epigenetic mechanisms controlling gene expression underlying growth and stress responses (Chinnusamy & Zhu 2009; Weng *et al.* 2014; Lee, Park & Seo 2017). There are numerous reports which suggested the role of histone acetylation and methylation (specifically H3K4me2 and H3K4me3) in transcriptional activation underlying stress responses (Saleh *et al.* 2008; Hu *et al.* 2015; Hu, Lu, Zhao & Zhou 2019; Ueda & Seki 2020). We, therefore, tested the histone acetylation and methylation at the promoters of HS genes in plants treated with and without glucose (+/− Glc). ChIP assay was performed using anti-histone 3 (K9+14+18+23+27) acetylation and anti-H3K4me2 antibodies and approximately 200 bp upsteram promoters of HS genes were analysed using ChIP-qPCR. Glucose alters the H3 acetylation (K9+14+18+23+27) and H3K4me2 at the promoters of HS genes (Figure 5B, Supplemental Figure S7). Presence of Glc showed more accumulation of these epigenetic marks at the promoters of HS genes encompassing HSEs than plants treated without glucose (Figure 5B, Supplemental Figure S7). We next asked whether TOR is required for glucose induced epigenetic changes at HS gene promoters. Five-day-old Arabidopsis *tor-es1* seedlings were transferred to estradiol (20µM) containing MS medium and kept for four days followed by starvation for 24h and 3h Glc treatment. Mock seedlings under Glc sufficiency showed increased accumulation of these histone acetylation and methylation marks at the HS gene promoters. On the contrary, glucose could not induce the accumulation of these epigenetic marks in seedlings treated with estradiol (Figure 5B, Supplemental Figure S7). Collectively, these results suggest the epigenetic mode of HS gene regulation by glucose-TOR signaling.

**Figure 5.**
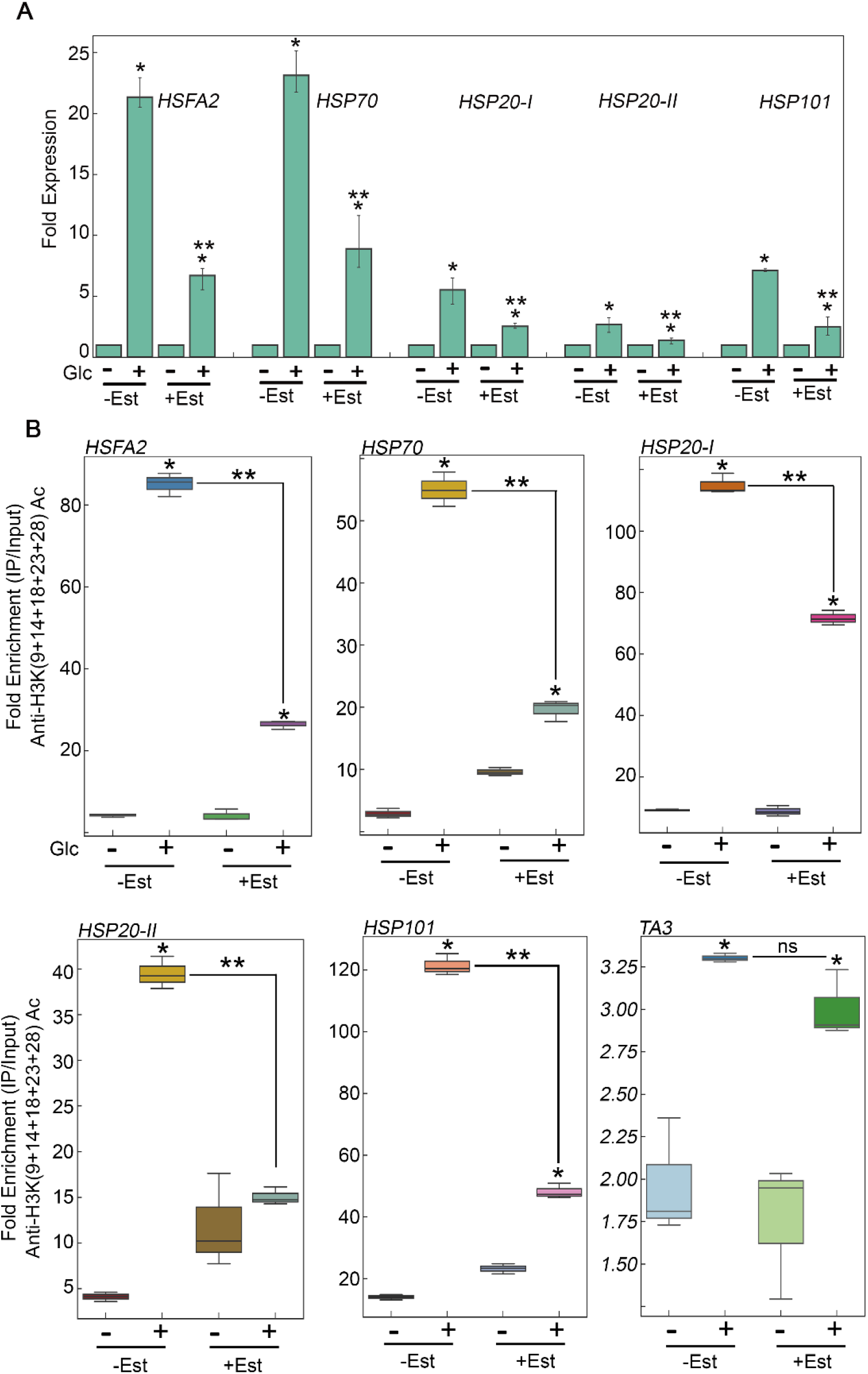
Glucose through TOR drives HS gene expression by incorporating histone H3 acetylation. **A**, RT-qPCR expression of HS genes in Arabidopsis *tor* estradiol inducible RNAi lines (*tor-es1*) under Glc lacking/sufficiency. Data shown are average of three biological replicates. **B**, ChIP-qPCR showing enrichment of histone H3K acetylation at the promoters of HS loci under Glc lacking/sufficiency. Promoter fragments containing cis-acting HSEs were immuno-precipitated using anti-H3K ac (9+14+18+ 23+27) antibody. Amount of immuno-precipitated promoter DNA was calculated by comparing samples treated without or with anti-Histone H3K (9+14+18+ 23+27) acetyl antibody. Ct values without and with antibody samples were normalized by input control. TA3 is a highly heterochromatinized DNA and was used as a negative control. In **A, B**, Five-day-old *tor-es1* seedlings were transferred to MS medium containing 20µM β-estradiol for four days. Following estradiol treatment, seedlings were subjected to 24h energy starvation in MS medium without Glc and then supplied with 3h Glc treatment. For mock treatment, seedlings were transferred in equal volume of DMSO (as used for estradiol) containing MS medium. Data shown are representative of one biological replicates. Experiments were repeated thrice with similar results. Error bars = SD (Student’s t test, P, 0.05; *control versus treatment; **Mock versus Estradiol).

### TOR coordinates with histone acetylation machinery to regulate sugar-mediated acetylation

Since Glc and TOR regulate HS genes expression by modulating epigenetic status, it was important to assess how Glc/TOR controls these responses. In mammals, TOR through phosphorylation activates a CREB-binding protein/p300-type histone acetyltransferase (Wan *et al.* 2017). In Arabidopsis, ortholog of p300 histone acetyltransferase HAC1 undergoes phosphorylation in response to sucrose (Van Leene *et al.* 2019). Moreover, HAC1 in combination with WRKY18/53 is recruited at the promoters of sugar responsive genes and regulate their acetylation under sugar sufficiency (Chen *et al.* 2019). To examine whether TOR mediated epigenetic changes at HS gene promoters are governed through HAC1, we assessed the expression network of Arabidopsis HAC1 with TOR and RAPTOR1. Plants gauge and adapt to diurnal light-dark cycle through transcriptional networks which facilitates plant’s response to external environment (Ferrari *et al.* 2019). We, therefore, assessed the comparative expression of *TOR* and *HAC1* in diurnal light-dark cycle transcriptome data obtained from public resources and observed that *TOR* concurs in expression with *HAC1* throughout diurnal cycle (Ferrari *et al.* 2019) (Supplemental Figure 8A). In addition, data obtained from Genevestigator (https://genevestigator.com/) and string (http://string-db.org/), we observed that *HAC1* shares huge similarity with *TOR* and *RAPTOR1* expression in anatomy, development and perturbations (Supplemental Figure 8B-D, Supplemental Figure 11). Next, to assess the HS hypersensitivity of *hac1* mutants, we examined the phenotype of Arabidopsis *hac1* in response to HS. Seven-day-old Arabidopsis Col-0 and *hac1* seedlings (*hac1-2* and *hac1-6*) were subjected to Glc free and Glc containing MS media for 24h followed by short term acquired thermotolerance (SAT) HS treatment. Col-0 and *hac1* mutants treated without Glc could not survive the lethal HS. However, Col-0 plants supplemented with Glc showed improved thermotolerance (showed higher fresh weight, increased chlorophyll [Chl] and higher lateral root numbers) (Figure 6A and 6B). Interestingly, *hac1* mutants showed reduced thermotolerance as observed by lesser fresh weight and Chl content and fewer lateral roots than Col-0 (Figure 6A and 6B). Next, we assessed the expression levels of HS genes underlying thermotolerance response in Col-0 and *hac1* using RT-qPCR. Alleles of *hac1* (*hac1-2*, *hac1-3* and *hac1-6*) revealed greater perturbation in HS gene expression following sugar and HS treatment than Col-0 seedlings (Figure 6C; Supplemental Figure S9A and S9B). To examine whether HAC1 regulates the expression of HS gene by affecting histone acetylation at HS responsive loci, we performed ChIP assay in Col-0 and *hac1* seedlings under Glc and HS treatment using anti-H3K (9+14+18+23+27) acetylation antibody coupled with qPCR. Indeed, *hac1* showed a greater reduction in the accumulation of histone acetylation at HS loci under Glc and HS treatment compared to Col-0 seedlings (Figure 6D, Supplemental Figure S9C). In addition, global transcriptome of *tor* DEGs with *hac1/5* obtained from public resources (Jin *et al.* 2018) identified 269 commonly down-regulated genes, with high enrichment of processes related to protein folding or heat shock responses (Supplemental Figure S10A-C). Furthermore, TOR and RAPTOR1 exhibited huge overlap of co-expression network with HAC1 with high enrichment of processes involved in chromatin remodelling, histone modification and gene transcriptional/posttranscriptional regulation (Supplemental Figure S10D-F). Collectively, these findings suggest that Glc-TOR signaling works in concert with the epigenetic machinery to regulate the expression of HS signaling genes, leading to thermotalerance.

**Figure 6.**
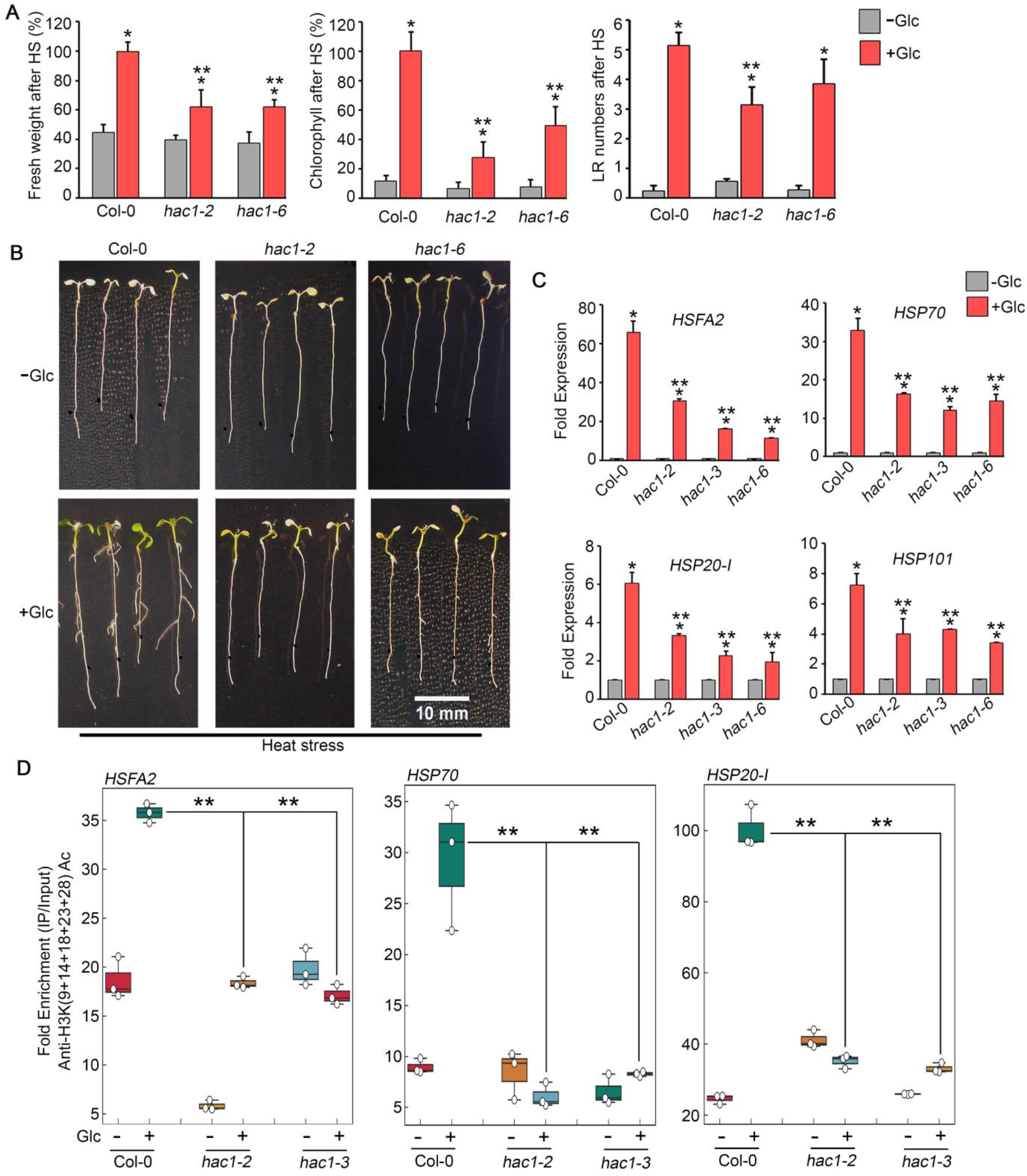
Glc-TOR mediated HS response in governed through HAC1. A, Percentage measurement of fresh weight, Chl content and LR numbers of Arabidopsis Col-0 and *hac1* mutants after HS. **B**, HS phenotype of Arabidopsis Col-0 and *hac1* seedlings primed with Glc. Five-day-old MS grown Arabidopsis seedlings treated without or with Glc and subjected to HS. HS was applied as 1h_37ºC, 2h_22ºC, 2.5h_45ºC followed by recovery at 3-4d_22ºC. Data shown are average of four (Fresh weight and Chlorophyll) and three (Lateral root number) biological replicates. **C**, RT-qPCR expression of HS genes in Col-0 and *hac1* mutants in response to Glc. **D**, ChIP-qPCR showing enrichment of histone H3K ac (9+14+18+ 23+27) at the promoters of HS gene encompassing HSEs in Col-0 and *hac1* mutants. Amount of immuno-precipitated promoter DNA was calculated by comparing samples treated without or with anti-Histone H3K (9+14+18+ 23+27) acetyl antibody. Ct values without and with antibody samples were normalized by input control. In **C**, **D**, seven-day-old MS grown Col-0 and *hac1* seedlings were subjected to 24h energy starvation in MS medium without Glc and then supplied with 3h Glc treatment. Data shown are representative of one biological replicates. Experiments were repeated thrice with similar results. Error bars = SD (Student’s t test, P, 0.05; *control versus treatment; **WT versus mutants).

### Glucose activated TOR is also required for maintaining short and long term thermomemory

Under natural environment, plants are constantly challenged by recurrent often irregular exposure of heat waves. Therefore, it is important to remember the environmental history as primed plants are more resistant to future stresses than naïve plants. It has already been proposed that Glc establishes and maintains the sustained expression of thermomemory-associated genes, and thereby causing thermomemory (Sharma *et al.* 2019). It is also necessary to test whether Arabidopsis TOR only controls acquired thermotolerance or to recall previous exposure to heat stress. To check this, we analysed the transcripts of thermomemory-associated genes in *tor35-7 RNAi* line. Seven-day-old Arabidopsis plants grown under moderate Glc (56mM) were treated with HS and recovered for various time points (3h, 24h and 48h). HS was applied as follows: 1h_37ºC, 90min_22ºC, 45min_45ºC and then recovered at 22ºC for 3h, 24h and 48h. Intriguingly, we observed that *tor35-7* RNAi showed reduced expression of thermomemory genes after extended recovery (24h and 48h) as compared to Col-0 (Figure 7A).

**Figure 7.**
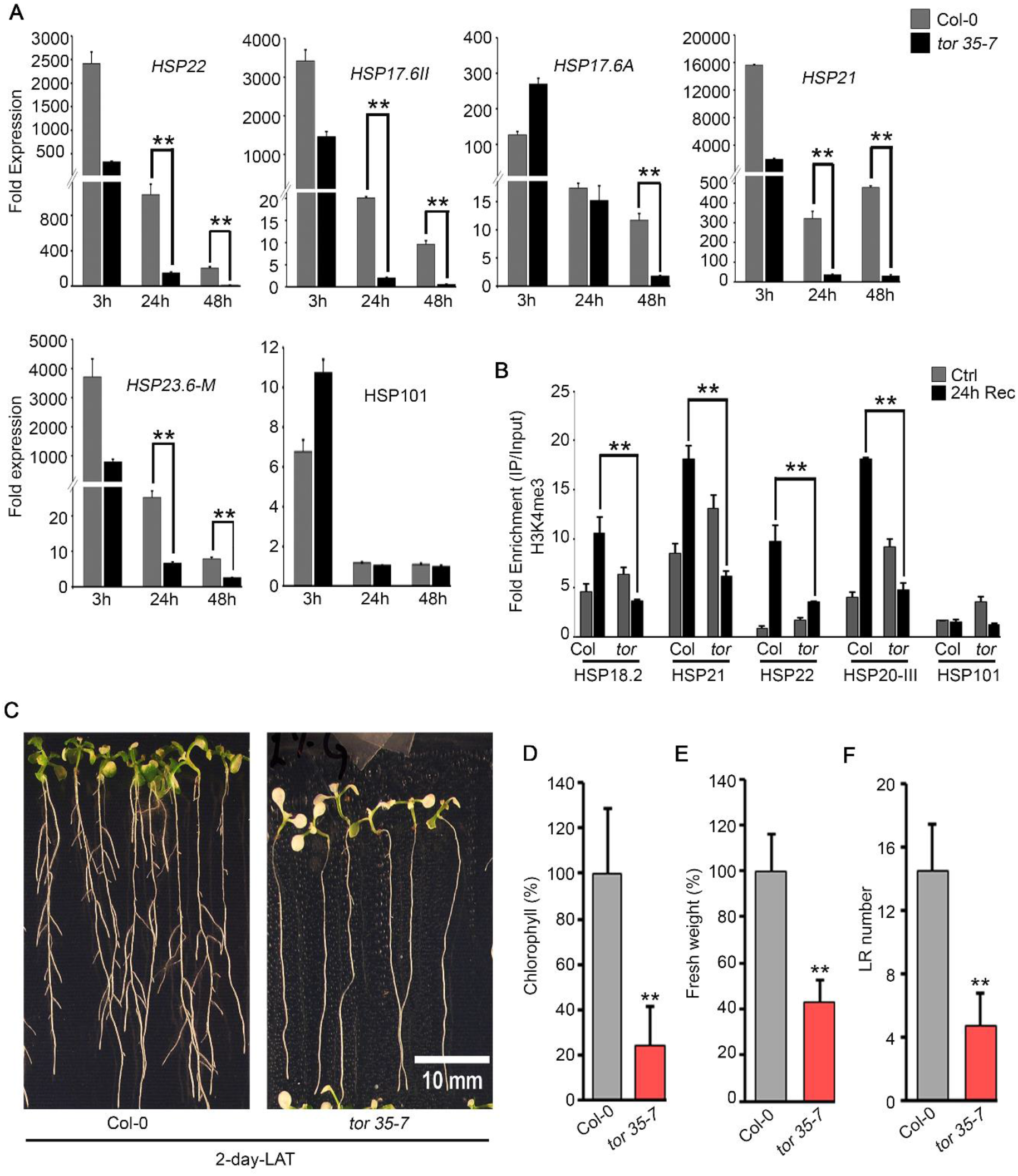
Glc regulated TOR is required in maintaining thermomemory. **A**, RT-qPCR expression of thermomemory related genes in Arabidopsis Col-0 and *tor 35-7 RNAi* seedlings. Arabidopsis Col-0 and *tor 35-7* plants were grown for seven days in presence of 1% Glc (56mM) under long-day conditions (16h light and 8h dark, 100 μM m-2 s-1 light intensity) treated without or with HS. HS was applied as 1h_37ºC, 90min_22ºC, 45min_45ºC. After HS, seedlings were recovered for various time points (3h, 24h and 48h) at 22ºC. Data shown are representative of four technical replicates among two biological replicates. Experiments were repeated twice times with similar results. **B**, ChIP-qPCR enrichment of thermomemory related genes in Col-0 and *tor 35-7* seedlings treated without or with HS. Following HS, seedlings were recovered for 24h at 22^º^C. Amount of immunoprecipitated promoter DNA was calculated by comparing samples treated without or with anti-Histone H3K4me3 antibody. C_T_ Values without and with anti-Histone H3K4me3 antibody were normalized by input control. Data shown are representative of one biological replicate. Experiments were repeated four times with similar results. **C**, Picture showing seedling survival of Arabidopsis Col-0 and *tor 35-7* RNAi after long term acquired thermotolerance. **D-F**, Percentage Chl, fresh weight and LR count of Arabidopsis Col-0 and *tor 35-7* RNAi after LAT HS. Arabidopsis Col-0 and *tor 35-7* plants were grown for seven days in presence of 1% Glc (56mM) under long-day conditions (16h light and 8h dark, 100 μM m-2 s-1 light intensity) and subjected to HS. HS was applied as 1h_37ºC, 2d_22ºC, 2.5h_45ºC and 3-4d_22ºC. Data shown are representative of four biological replicates. Experiments were repeated four times with similar results.

**Figure 8.**
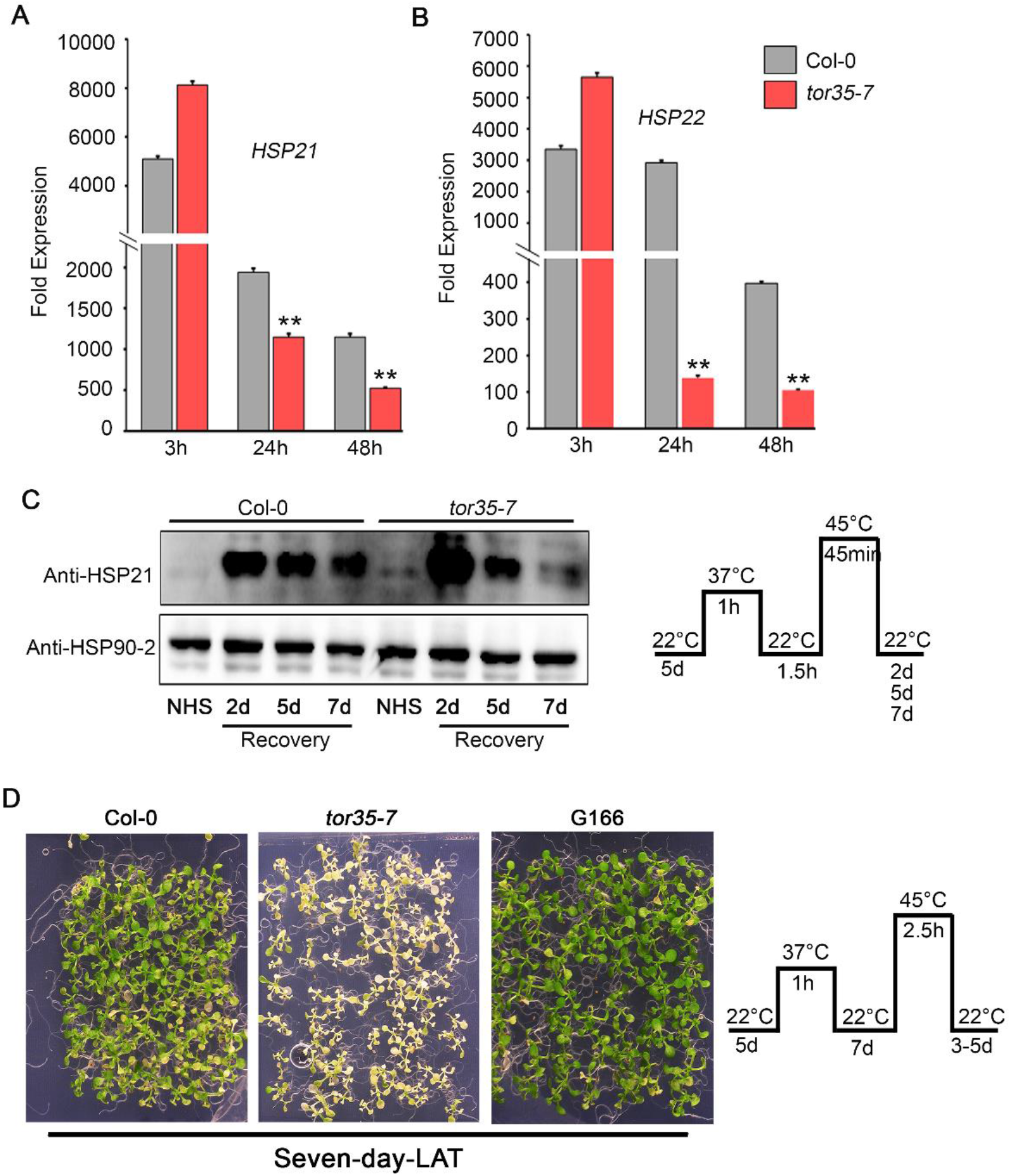
TOR controls the duration of thermomemory. **A, B**, RT-qPCR expression of thermomemory related genes in Arabidopsis Col-0 and *tor 35-7 RNAi* seedlings after LAT HS. Arabidopsis Col-0 and *tor 35-7* plants were grown for seven days in presence of 1% Glc (56mM) under long-day conditions (16h light and 8h dark, 100 μM m-2 s-1 light intensity) treated without or with HS. HS was applied as 1h_37ºC, 2d_22ºC, 45min_45ºC. After HS, seedlings were recovered for various time points (3h, 24h and 48h) at 22ºC. Data shown are representative of one biological replicate. Experiments were repeated twice with similar results. **C**, Immunoblot detection of thermomemory-associated HSP21 protein in Arabidopsis Col-0 and tor 35-7 seedlings after 2-7d of HS recovery. Arabidopsis plants were grown for five days in MS medium containing 1%Glc (56mM) and subjected to HS as 1h_37ºC, 1.5h_22ºC, 45min_45ºC. Following HS, seedlings were recovered for 2d, 5d and 7d at 22ºC. HSP21 protein was detected using anti-HSP21 specific antibody and HSP90 was detected using Anti-HSP90-2 specific antibody and used as a loading control. Data shown are representative of one biological replicate. Experiment was repeated thrice with similar result. **D**, Phenotype of Arabidopsis Col-0 and tor 35-7 plants after seven-day of LAT HS. Arabidopsis Col-0 and *tor 35-7* plants were grown for five days in MS medium containing 1% Glc (56mM) and subjected to HS. HS was applied as 1h_37ºC, 7d_22ºC, 2.5h_45ºC and 3-5d_22ºC. Picture shown are representative of one biological replicate. Experiment was repeated twice with similar results.

Chromatin structures are crucial factors which remember past experience of stress and show quick and heightened response to future stresses (Struhl & Segal 2013; Zentner & Henikoff 2013). Enthralling evidences revealed the critical roles of histone methylation specifically H3K4me3 in chromatin based transcriptional memory (Ding *et al.* 2012b a; Lämke *et al.* 2016; Sharma *et al.* 2019). We, therefore, analysed the H3K4me3 epigenetic marks at the promoters of thermomemory associated loci after 24h of HS recovery in Col-0 and *tor35-7*. Col-0 plants showed higher accumulation of H3K4me3 at the promoters of thermomemory loci whereas *tor35-7 RNAi* plants failed to accumulate these marks (Figure 7B). Next, we asked whether TOR is only able to remember short duration of HS (SAT) or is involved in long term acquired thermotolerance (LAT). To this end, we examined the phenotype of Arabidopsis Col-0 and *tor35-7* after LAT HS. Seven-day-old Arabidopsis plants grown under moderate Glc (56mM) conditions were subjected to HS. HS was applied as follows: 1h_37ºC, 2d_22ºC, 2.5h_45ºC, 3-4d_22ºC. Arabidopsis Col-0 plants showed higher seedling survival whereas *tor35-7* seedlings failed to recover after LAT (Figure 7C-F). In addition, we checked the phenotype of TOR OE lines after LAT and priming and unpriming HS. Remarkably, TOR OEs showed higher seedling survival in comparison with Col-0 and *tor35-7* after LAT HS (Supplemental Figure S12A-C). In addition, TOR OEs survived better (as shown by robust phenotype) than Col-0 under priming HS, whereas no major changes were observed in Arabidopsis Col-0 and G166 seedlings without priming (Supplemental Figure S13A-D). However, G548 showed better growth recovery even after unpriming HS than Col-0 plants (Supplemental Figure S13A-D). Next, to assess whether high temperature affects TOR promoter activity, we examined transgenic line carrying TOR promoter-reporter fusion construct. In line, *pTOR::GUS* activity was largely restricted to shoot apical meristem (SAM) (Menand *et al.* 2002)). The plants of *pTOR::GUS* treated with Glc and recovered for 24h after HS showed strong reporter activity in the zones of primary meristem and at the leaf base (Supplemental Figure S14).

Furthermore, we also checked expression of thermomemory associated genes in *e2fa-1*. Similar to *tor35-7*, *e2fa-1* plants did not show sustained expression of memory genes than WT (Supplemental Figure S15). We next checked the long-term acquired thermotolerance (LAT) in Arabidopsis Col-0 and *e2fa-1* plants. Arabidopsis *e2fa-1* showed reduced leaf greening and Chl than Col-0 supporting the role of TOR/E2Fa signaling in providing thermomemory (Supplemental Figure S16A and S16B). Since *e2fa-1* showed reduced memory gene expression, we were interested to know whether E2Fa binds directly to thermomemory gene promoters to activate their expression. To this end, we performed ChIP assay in Arabidopsis Col-0 plants using anti-E2Fa antibody. Interestingly, E2Fa did not bind directly to these gene promoters (Supplemental Figure S16C). We also checked binding of E2Fa to its known targets such as MCM5 and ATXR6 as positive control (Raynaud *et al.* 2006; Xiong *et al.* 2013). Collectively, these results suggest that glucose involves TOR-E2Fa signaling in sustaining thermomemory associated gene expression to provide thermotolerance.

### Arabidopsis TOR controls the length of the transcriptional memory persistence

Induction of key thermomemory-related genes and their sustained expression is crucial and considered as potential factor in establishing thermomemory (Lämke *et al.* 2016; Sharma *et al.* 2019). We, therefore, looked at the expression of thermomemory-associated genes in Col-0 and *tor35-7* after extending intervening recovery time from two hours (SAT) to two days (LAT). Arabidopsis *tor35-7* showed decreased sustenance of thermomemory-related gene expression following 24h and 48h of recovery than Col-0 plants (Figure 8A and 8B). Furthermore, we examined the persistence of thermomemory-associated HSP21 protein in Col-0 and *tor35-7* after extending memory phase from two to seven days. Unexpectedly, *tor35-7* showed significantly higher levels of HSP21 protein after two days of HS recovery than Col-0 plants (Figure 8C). However, degradation of HSP21 was significantly faster after five and seven days of HS recovery in *tor35-7* (Figure 8C). Next, to test how long TOR controls the length of transcriptional memory, we primed Col-0, TOR OE and *tor35-7* plants with mild HS (37°C_3h) and extended the duration of intervening non-stress phase (22°C) up to seven days and then subjected to lethal HS (45°C_2.5h). Fascinatingly, TOR OE survived and recovered well from the deleterious HS. In contrast, *tor35-7* plants forgot their HS memory and showed greater sensitivity to HS than Col-0 (Figure 8D). Collectively, these several lines of evidence suggest the pivotal role of Arabidopsis TOR in establishment and maintenance of thermomemory response.

### TOR co-expression and transcriptome share huge overlap with transcriptome of chromatin remodeler BRAHMA

Since TOR regulates the epigenetics of non-memory and thermomemory-associated gene loci, it was apparent to assess the relationship between TOR and chromatin machinery. We analysed the co-expression network of TOR using *ATTED-II* database (http://atted.jp). We observed that TOR co-expresses with CHR2/ BRM and SYD (CHR3; Figure 9A). Moreover, E2Fa was found to be an interactor of BRM, SYD and SWI3C chromatin remodelers (Efroni *et al.* 2013) (https://thebiogrid.org; Supplemental Figure 17A). Previously, Brzezinka et al., 2016 showed that *brm-5* display reduced thermomemory. To examine whether TOR triggered thermomemory is mediated through BRM, we analysed the TOR co-expression network with BRM. Out of 2000 TOR co-expressed genes, 1474 (74%) genes were overlapping with BRM and 1236 (62%) genes were overlapping between TOR, BRM and CHR3 (Figure 9B; Supplemental Table S4A). Similarly, 966 genes were commonly co-expressed between TOR, BRM and RAPTOR1B (Supplemental Figure 18A-C; Supplemental Table S4B). BRM regulates target gene expression through the concerted functions of REF6 on a specific genomic sequence CTCTGYTY (Lu, Cui, Zhang, Jenuwein & Cao 2011; Li *et al.* 2016). GO of these common genes exhibited up-regulation of processes involved in chromatin remodelling/organization, histone posttranslational modifications and transcription co-regulator activity (Figure 9B; Supplemental Figure S18A-B; and S19A-B). Next, to test whether BRM and TOR regulates similar targets, we tested the DNA motifs recognised by BRM/REF6 in the genes co-expressed with TOR, *tor_DEG* and E2F motifs containing genes. Interestingly, in all three categories, we found approximately one third genes containing BRM/REF6 binding CTCTGYTY motifs in their 1000 bp upstream and 200 bp downstream promoters (Figure 9C; Supplemental Table S5). Next, we checked the phenotype of *brm-5* and *ref6-2* for seven-day-LAT. Intriguingly, *brm-5* and *ref6-2* could not remember long periods of HS memory and died as compared to Col-0 plants (Figure 9D and 9E). Further, we overlap *tor_DEG* with BRM and REF6 targets. Similar to co-expression, *tor_DEG* shares huge overlap with BRM and REF6 targets. Out of 4979 *tor_DEG*, 1392 and 611 genes were found to be overlapping with BRM and REF6 targets, respectively (Figure 9F; Supplemental Table S4C). GO enrichment of these common genes showed categories with phosphorelay signal transduction, abiotic stress response, transcriptional regulatory DNA binding and transcriptional regulator activity (Figure 9F-H). In addition, we also compared the genes containing E2F motifs with BRM/REF6 targets and found GO enrichment exactly similar to what we observed in *tor_DEG* comparison with BRM and REF6 (Supplemental Figure S19C-E; Supplemental Table S4D). In addition, only 173 (8.65%) genes were found to be overlapping between TOR, BRM and E2Fa co-expression (Supplemental Figure S18D-F; Supplemental Table S4E). Among 2000 co-expressed genes of TOR, RAPTOR, E2Fa and BRM, we fetched the genes involved in chromatin remodelling and posttranslational modifications. TOR and RAPTOR exhibited a co-expression pattern similar to BRM for most of these chromatin regulators (Supplemental Table S6). Moreover, phospho-proteome data of sucrose sufficiency/AZD-8055 treatment identified 4,988 sugar induced phospho-peptides in 2,119 proteins (Van Leene *et al.* 2019). Among these phospho-peptides, sucrose treatment induced phosphorylation in 26 proteins (containing 83 phospho-peptides) involved in chromatin machinery including BRM (15 peptides), SYD (8 peptides) and REF6 (7 peptides; Supplemental Figure S17B). Collectively, these extensive network analyses of TOR with BRM indicate that TOR may act in combination with BRM to regulate downstream processes, including thermotolerance/thermomemory.

**Figure 9.**
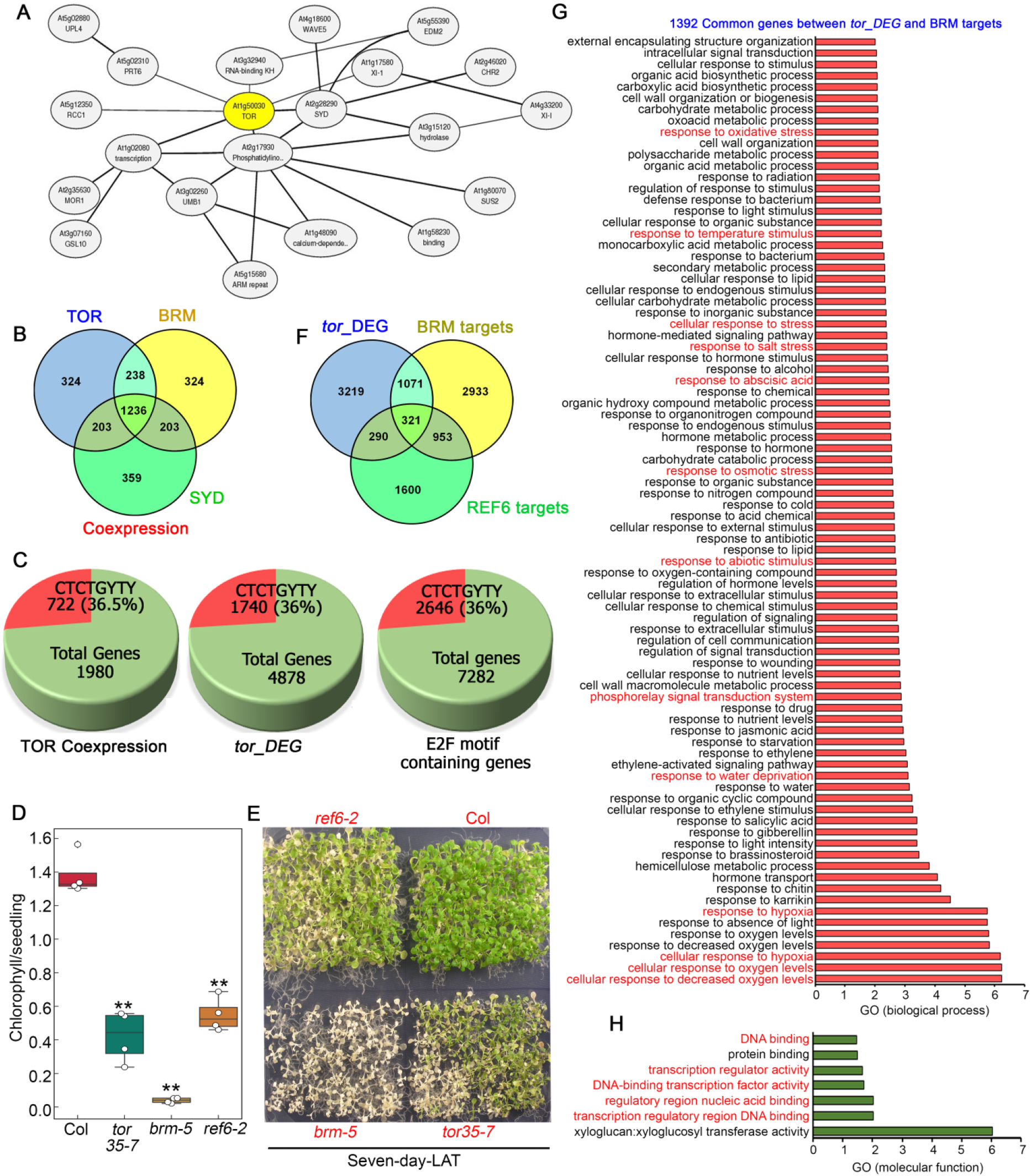
TOR co-expresses with chromatin remodeler BRM and overlaps common downstream targets. **A**, Network showing co-expression data of TOR with BRM and SYD (CHR3). Co-expression network was extracted from ATTED-II. **B**, Venn diagram showing overlap of TOR co-expressing genes with BRM and SYD. **C**, Pictures showing enrichment of BRM-REF6 recognising genomic motif CTCTGYTY in TOR co-expressing genes, tor_DEG and in genes containing E2Fa motifs in their promoters. 1000 bp upstream and 200 bp downstream Promoters relative to TSS was considered for promoter analysis. **D, E** Chlorophyll measurement and phenotype of Arabidopsis Col-0, *tor35-7*, *brm-5* and *ref6-2* plants after seven-day of LAT HS. Arabidopsis Col-0, *tor35-7*, *brm-5* and *ref6-2* plants were grown for five days in MS medium and subjected to HS. HS was applied as 3h_37ºC, 7d_22ºC, 2.5h_45ºC and 3-5d_22ºC. Picture shown are representative of one biological replicate. Experiment was repeated thrice with similar results. **F**, Venn diagram showing overlap of genes differentially regulated by TOR with BRM and REF6 targets. **G, H**, GO fold enrichment of biological process and molecular function of 1392 genes commonly regulated by *tor_DEG* and BRM targets. Panther 15.0 tool was used to analyse GO fold enrichment using Bonferroni correction and Fisher’s exact test type.

## DISCUSSION

Due to their stable nature, plants cannot move to more favourable locations and therefore must adapt to stressful temperatures through an inherent potential called basal or acquired thermotolerance. Furthermore, unlike animals, plants do not have the brain or antibodies to recall previous exposure to adverse conditions, for example illness or stress. Recent reports provide enthralling evidences of “epigenetic memory” which requires changes through alteration in nucleosome positioning, chromatin modification, posttranslational histone modifications and RNA pol-II stalling (Ding *et al.* 2012a; Sani *et al.* 2013; Zentner & Henikoff 2013; Tessarz & Kouzarides 2014; Venkatesh & Workman 2015; Brzezinka *et al.* 2016; Lämke *et al.* 2016; Liu *et al.* 2019; Sharma *et al.* 2019). Mechanistic understanding of transcriptional memory of recurrent stresses is most advanced in yeast (Brickner *et al.* 2007; Tan-Wong, Wijayatilake & Proudfoot 2009; Light, Brickner, Brand & Brickner 2010). In plants, transcriptional memory is underexplored and mostly described for plant defence responses (Bruce, Matthes, Napier & Pickett 2007; Conrath 2011; Jaskiewicz, Conrath & Peterhälnsel 2011; Avramova 2015). A number of reports exhibited involvement of epigenetic regulators in plant stress responses (Hu *et al.* 2015; Liu, Feng, Li & He 2015), however, few examples of transcriptional memory have been known in case of plant memory to abiotic stresses (Ding *et al.* 2012a; Lämke *et al.* 2016; Lämke & Bäurle 2017; Liu *et al.* 2019; Sharma *et al.* 2019). Through a comprehensive chemical, genetic, transcriptome and epigenomic analysis, our results provide the pivotal role of Glc-TOR signaling in HS adaptation and maintaining thermomemory. Previously, numerous researchers have identified the involvement of TOR in stress resilience and adaptation (Mahfouz, Kim, Delauney & Verma 2006; Deprost *et al.* 2007; Bakshi *et al.* 2017; Dong *et al.* 2017; Wang *et al.* 2018; Fu, Wang & Xiong 2020). Recently, Sharma et al., (2019) revealed that Arabidopsis TOR controls transcription of many heat shock protein (HSP) genes including HSFA2 and HSP70 (Sharma *et al.* 2019). However, the underlying mechanism how TOR controls thermotolerance response is not well understood. In line with the previous observation, transcriptome study of Arabidopsis seedlings under Glc lacking/sufficiency demonstrated that Glc largely affects the expression of HS-related genes. At the whole genome level, 1124 genes were co-regulated by −Glc/+HS and +Glc/+HS. As expected, several metabolic processes related to primary and secondary metabolism, microtubules disassembly, sulphur compound biosynthetic processes and biotic stimulus were severely down-regulated whereas genes involved in HS protection such as HS responsive molecular chaperones, de novo protein folding, HS acclimation and de novo post-translational protein folding were up-regulated (Supplemental Figure S2A and S2B). Interestingly, apart from common genes regulated by −Glc/+HS and +Glc/+HS, presence of Glc exclusively regulates 1408 genes. GO enrichment of these genes showed high enrichment of protein deneddylation process (Figure 2E). Protein neddylation and deneddylation upon stress is required for the assembly and disassembly of stress granules (SG), respectively (Jayabalan *et al.* 2016). Under stress, organisms show inhibition in protein synthesis, blocks translation initiation and triggers polysome disassembly and sequestration of translation stalled mRNA into aggregates called SG for storage (Chantarachot & Bailey-Serres 2018). SGs prevent the degradation of mRNA and store them which can reversibly be engaged into the translation once stress has removed. Recently, Merret et al., (2017) showed that mutants of *HSP101* showed perturbation in translational recovery and SG disassembly following HS (Merret *et al.* 2017). In agreement, our data indicate that Glc signaling may regulate SG disassembly through expression of genes underlying deneddylation which are crucial for stress recovery. In addition, Glc through TOR promotes the transcriptional regulation of *HSP101* which is crucial for SGs disassembly and may therefore provide an adaptive strategy of plant survival under deleterious HS. Transcriptome reprogramming of HS genes under Glc sufficiency exhibited major synergistic interaction (310 down- and 250 simultaneously up-regulated) with TOR regulated genes suggesting that Glc governs transcriptome reprogramming of HS genes mainly through TOR pathway.

Another important feature of HS response was to study the transcriptome of genes induced/suppressed when the HS signal was absent, i.e. HS recovery. Recently, our group has shown that Glc maintains the sustained expression of HS genes, thereby regulates thermomemory (Sharma *et al.* 2019). We found that a huge number of genes (1332; 46%) demonstrated their sustained expression when HS stressed plants were recovered under Glc sufficiency. These include GO processes related to DNA unwinding, geometrical and conformational changes, DNA replication, ribosome biogenesis and cell wall modifications, which may be required for new cell divisions following stress removal (Supplemental Figure S5B and S5C). In addition, HS recovery under Glc specifically induced GO processes related to histone/protein methylation which perhaps required to facilitate stress memory response (Figure 3F; Supplemental Figure S5B). In contrast, expression profile of genes was completely vanished (only 28 genes; 1.2%) when plants were recovered without Glc. Interestingly, comparison of the HS recovery transcriptome with TOR revealed that Glc sustenance of HS recovery genes was primarily governed through TOR. Earlier reports have shown that Glc regulates transcriptome reprogramming of myriad set of genes via E2Fa transcription factor (Xiong *et al.* 2013; Li *et al.* 2017). We observed that E2Fa recruited at HSFA1 and HSFA2 upstream genomic regions and positively regulates their expression.

Numerous reports revealed that dynamic nature of chromatin structure is a crucial determinant of gene expression (Berr *et al.* 2009; Tamada, Yun, Woo & Amasino 2009; Kumar & Wigge 2010; Brzezinka *et al.* 2016). Our study provides strong evidence supporting that Glc-TOR signaling regulates HS transcripts by accumulation of histone acetylation marks at the promoters of HS loci. We observed a strong defect of *tor* on Glc promoted HS expression, demonstrating that Glc induction of HS transcriptome is majorly regulated through TOR. Lately, an interactome and phosphoproteome study of sugar and TOR identified CBP/p300 protein HAC1 as plausible TOR target (Van Leene *et al.* 2019). In our study, we observed that *hac1* showed hyper-sensitivity to HS with a strong defect in Glc promoted accumulation of H3K acetylation, leading to reduced expression of the HS gene. Collectively, these results demonstrate that Glc-TOR acts co-ordinately with histone acetylation machinery to regulate genes underlying thermotolerance response.

Plant stress research has largely been focussed on basal or acquired tolerance to certain levels of stresses. However, the impact of environmental history on the plant performance to recurrent and irregular stresses has been under-researched. Plant memory to recurrent stresses can be described as non-epigenetic such as RNA metabolism (e.g., decay or stabilization), protein turnover and epigenetic mechanisms which involve lasting changes in chromatin (Charng 2006; Ding *et al.* 2012a; Brzezinka *et al.* 2016; Crisp, Ganguly, Eichten, Borevitz & Pogson 2016; Sedaghatmehr, Mueller-Roeber & Balazadeh 2016). TOR, as being an important regulatory gene governs numerous biological processes ranging from embryo development to senescence and stress responses (Sheen 2014; Fu *et al.* 2020). However, the involvement of TOR in plant response to recurrent or repeated HS is hitherto unexplored. Here, we demonstrated that TOR modulates the dynamics of chromatin status in repetitively facing HS. Numerous reports revealed preservation of chromatin modification specifically H3K4me3 at trained genes in response to variety of environmental stresses (Tsuji, Saika, Tsutsumi, Hirai & Nakazono 2006; Sokol, Kwiatkowska, Jerzmanowski & Prymakowska-Bosak 2007; Ding *et al.* 2012a; Singh *et al.* 2014; Lämke *et al.* 2016). Interestingly, Arabidopsis plants exposed to HS demonstrated increased accumulation of histone H3K4me3 marks, leading to more open chromatin in the promoters of thermomemory-associated genes which was greatly suppressed in plants defective in TOR. This increased enrichment of H3K4me3 at thermomemory associated gene promoters correlated with the sustained expression over a period of days, leading to increased thermomemory. We have also shown that Arabidopsis TOR not only demonstrated thermomemory response to SAT, LAT responses were also governed by TOR. Arabidopsis WT plants exposed to LAT HS showed increased sustenance of thermomemory associated *HSP21* and *HSP22* gene transcripts which was compromised in *tor35-7* plants. Moreover, we observed that *tor35-7* displayed higher degradation of HSP21, thereby causing greater sensitivity to repeated HS. Given the importance of chromatin remodelling in stress and memory response, we observed significant overlap of TOR co-expression and DEGs with BRM. In addition, we found the significant presence of BRM cis-acting motifs in the genomic loci of TOR co-expression, *tor_DEG* and E2F motifs containing genes.

In conclusion, these results suggest a central role of Glc-TOR signaling along with transcriptional and chromatin machinery which are thought to act in concert to control thermotolerance/thermomemory (Figure 10). This work identified the crucial function of TOR in shaping the environmental history of plants and provides a mechanistic framework to confront ever challenging and fluctuating environment. This approach would be exploited to improve temperature tolerant crop varieties in order to feed ever growing human population.

**Figure 10.**
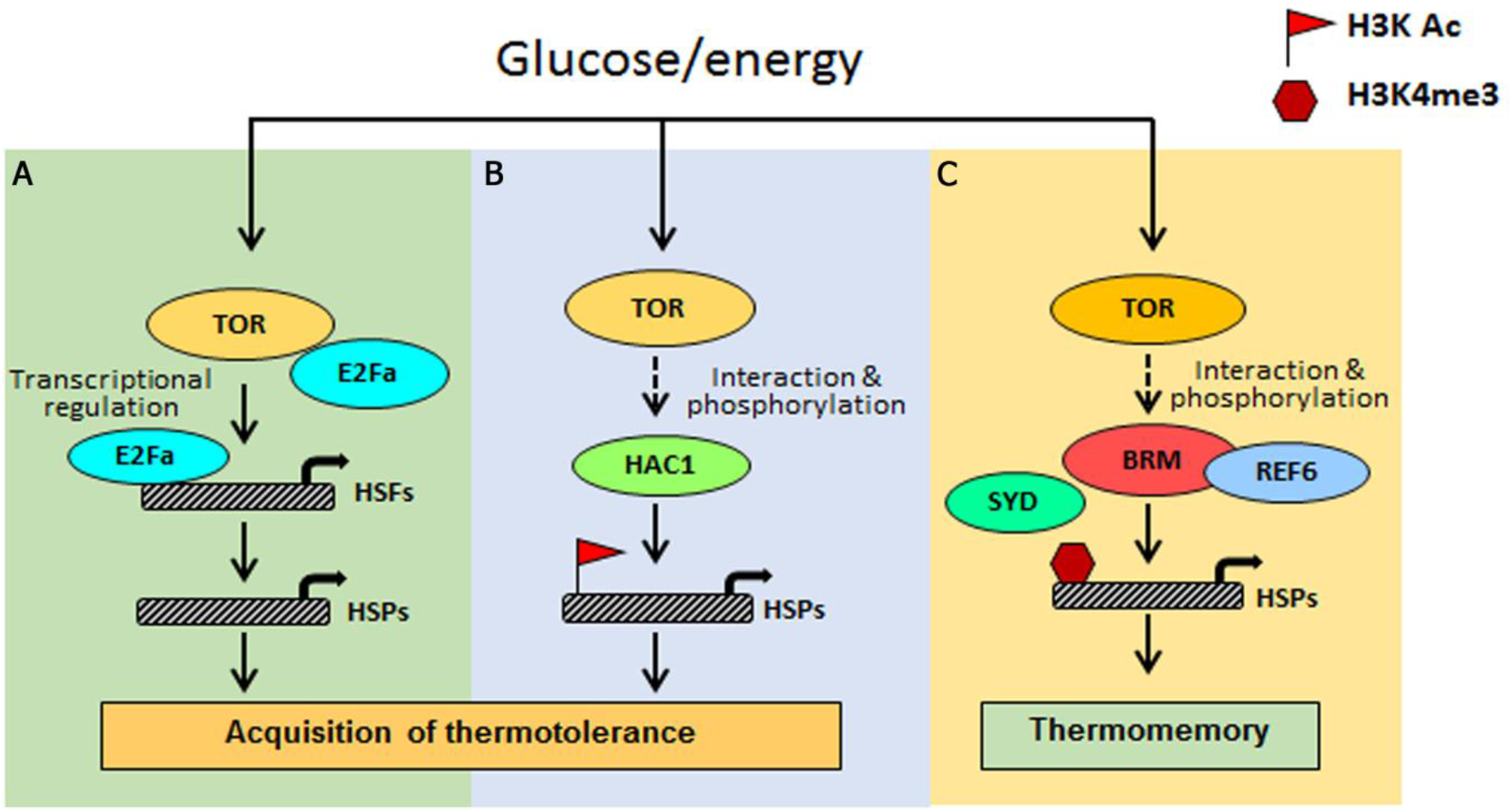
Proposed model shows how Glc-TOR signalling through distinct mechanisms regulates thermotolerance/thermomemory. **A**, Arabidopsis TOR regulated E2Fa transcription factor recruited at the promoters of HSFA1 and HSFA2 genes to control their expression which in turn regulate HS gene expression. **B**, Glc through TOR promotes recruitment of histone H3 acetylation marks at the promoters of HS genes to induce their transcription, leading to thermotolerance. This Glc-TOR mediated histone acetylation is facilitated through HAC1. **C**, Sugar also controls interaction and phosphorylation of several chromatin proteins including BRM, SYD and REF6. TOR co-expresses with BRM & SYD and exhibits huge overlap with BRM target genes. TOR promotes the accumulation of histone methylation marks at the promoters of thermomemory associated gene promoters, and therefore regulates thermomemory.

## ACKNOWLEDGEMENTS

We are grateful to Prof. Lieven De Veylder, VIB Department of Plant Systems Biology, Ghent University, Belgium for anti-E2Fa serum. We thank Prof. Christian Meyer, French National Institute for Agricultural Research, INRA, Institut Jean-Pierre Bourgin for providing *tor 35-7* RNAi lines. We thank Benoît menand, CNRS research scientist, Plant genetics and biophysics laboratory (IBEB / UMR7265 CEA-CNRS-AMU), Luminy University Campus (Marseille, 13) for providing pTOR::GUS seeds. We thank Prof. Susan I. Gibson, Department of Plant Biology, Microbial and Plant Genomics Institute, University of Minnesota, Saint Paul, MN, USA for providing HAC1 mutant seeds. We thank Central Instrumental Facility, NIPGR for all the required assistance. We thank Dr. Sandhya Verma and Ms. Rekha Agrawal for advice, valuable discussions and critical reading of the manuscript. We thank Dr. Jitendra K. Thakur for providing laboratory space for performing few experiments. This work was supported by core grant of National Institute of Plant Genome Research, New Delhi, India to A.L. M.S., received fellowship from Department of Biotechnology (DBT) and Science and Engineering Research Board (SERB) with Grant Number: EMR/2016/006486. MJK acknowledges Department of Science and Technology (INSPIRE Faculty Programme Grant IFA18-LSPA110). M.S. received fellowship from University grant commission, P.A., and S.K.M. received fellowship from DBT. Promoter motif analysis, Diurnal and Genevestigator analysis were performed using funds in the DBT Grant Number-BT/PR22334/BID/7/786/2016. The authors are thankful to DBT-eLibrary Consortium (DeLCON) for providing access to e-resources.

## AUTHOR CONTRIBUTIONS

M.S. and A.L. conceived and designed the experiments. M.S. performed most of the experiments. M.S. and B.N.S. performed immuno-blotting experiment. M.S. and M.J.K. performed Microarray experiment. M.S., B.N.S., M.S. and P.A. performed physiology and RT-qPCR experiments. M.S., S.K.M. and G.Y. performed promoter motif analysis, Diurnal and Genevestigator analysis. M.S. and A.L. analysed the data. M.S wrote the manuscript. A.L. supervised and complemented the writing.

## Supplemental data

The following supplementary data are available:

**Supplemental Figure S1. Photosynthesis generated Glc could not provide thermotolerance under high light intensity when blocked with Glc analogue 2-deoxy-glucose. A**, Phenotype showing effect of Glc analogue 2-DG on thermotolerance response. **B-D**, Percentage fresh weight, chlorophyll content and lateral root numbers of Arabidopsis Col-0 seedlings after treatment with high light and heat stress in presence of Glc alone and Glc along with its analogue 2-DG. Five-day-old MS grown Arabidopsis Col-0 seedlings under standard light intensity were transferred to Glc (56mM) containing MS media without or with Glc analogue 2-DG (2.5mM). After transfer, seedlings were transferred to high light intensity (approximately 12000 LUX) for 24h and then subjected to heat stress at 1h_37ºC, 2h_22ºC, 2.5h_45ºC and 3-7d_22ºC. Data shown are average of three biological replicates. Experiment was repeated thrice with similar results. Error bars represent SD (Student’s t test, P < 0.05; **, Mock vs inhibitor).

**Supplemental Figure S2. Gene ontology of common genes between –Glc/+HS and +Glc/+HS exhibits up-regulation of genes involved in protein folding. A, B** GO term enrichment of biological process synergistically down-and up-regulated by both –Glc/+HS and +Glc/+HS. Top twenty GO categories were included based on their fold enrichment. Panther 15.0 tool was used to analyse GO fold enrichment using Bonferroni correction and Fisher’s exact test type.

**Supplemental Figure S3. Glc regulates HS transcriptome largely through TOR pathway. A**, GO biological process of genes exclusively down-regulated by –Glc/+HS. **B, C**, GO biological process of genes commonly down- and up-regulated by +Glc/+HS and public available Glc-TOR microarray genes (Xiong *et al.* 2013). Top twenty GO categories were included based on their fold enrichment. Panther 15.0 tool was used to analyse GO fold enrichment using Bonferroni correction and Fisher’s exact test type.

**Supplemental Figure S4. HS transcriptome induced in absence of Glc exhibits large antagonistic interaction with Glc-TOR target genes. A, B**, GO biological process of genes antagonistically (down-by Glc-TOR and up-regulated by –Glc/+HS; up-by Glc-TOR and down-regulated by –Glc/+HS) regulated by Glc-TOR and –Glc/+HS. **C, D**, GO biological process enrichment of genes synergistically (up- and down-regulated by both Glc-TOR and –Glc/+HS) regulated by Glc-TOR and –Glc/+HS. Top twenty GO categories were included based on their fold enrichment. Panther 15.0 tool was used to analyse GO fold enrichment using Bonferroni correction and Fisher’s exact test type.

**Supplemental Figure S5. Presence of Glc regulates sustenance of HS transcriptome for HS recovery. A**, GO biological process of genes commonly down-regulated by +Glc/+HS and +Glc/+HS Rec. **B, C**, GO biological process of genes exclusively up- and down-regulated by +Glc/+HS Rec in comparison with –Glc/+HS Rec. Top twenty GO categories were included based on their fold enrichment. Panther 15.0 tool was used to analyse GO fold enrichment using Bonferroni correction and Fisher’s exact test type.

**Supplemental Figure S6. HS recovery under Glc induces processes required for growth/HS recovery in a TOR dependent manner. A**, GO biological process of genes commonly up-regulated by +Glc/+HS Rec and TOR. **B**, GO biological process of genes commonly down-regulated by +Glc/+HS Rec and TOR. Top twenty GO categories were included based on their fold enrichment. Panther 15.0 tool was used to analyse GO fold enrichment using Bonferroni correction and Fisher’s exact test type.

**Supplemental Figure S7. Glc through TOR induces enrichment of histone H3K4me2 marks at the promoters of HS genes.** ChIP-qPCR showing enrichment of histone H3K4me2 at the promoters of HS loci under Glc lacking/sufficiency. Promoter fragments containing cis-acting HSEs were immuno-precipitated using anti-H3K4me2 antibody. Amount of immuno-precipitated promoter DNA was calculated by comparing samples treated without or with anti-H3K4me2 antibody. Ct values without and with antibody samples were normalized by input control. TA3 is a highly heterochromatinized DNA and was used as a negative control. Five-day-old *tor-es1* seedlings were transferred to MS medium containing 20µM β-estradiol for four days. Following estradiol treatment, seedlings were subjected to 24h energy starvation in MS medium without Glc and then supplied with 3h Glc treatment. For mock treatment, seedlings were transferred in equal volume of DMSO (as used for estradiol) containing MS medium. Data shown are representative of one biological replicates. Experiments were repeated twice with similar results. Error bars = SD (Student’s t test, P, 0.05; *control versus treatment; **Mock versus Estradiol).

**Supplemental Figure S8. Diurnal light-dark cycle and Genevestigator transcriptome comparison shows similarity in expression between TOR, RAPTOR1, HAC1 and BRM. A**, Heat map showing expression profiling of TOR, RAPTOR1, HAC1 and BRM in diurnal light-dark transcriptome data obtained from public resources (Ferrari *et al.* 2019). **B-D**, Heat maps showing expression profiling of TOR, RAPTOR1, HAC1 and BRM in anatomy, development and perturbations data obtained from Genevestigator.

**Supplemental Figure S9.***hac1* **mutant demonstrates less accumulation of H3K acetylation, leading to perturbation in HS induced gene expression. A, B**, RT-qPCR expression of HS genes in Col-0 and *hac1* mutants in response to HS. **C**, ChIP-qPCR showing enrichment of histone H3K Ac (9+14+18+ 23+27) at the promoters of HS gene encompassing HSEs in Col-0 and *hac1-2* mutant. Amount of immuno-precipitated promoter DNA was calculated by comparing samples treated without or with anti-Histone H3K (9+14+18+ 23+27) acetyl antibody. Ct values without and with antibody samples were normalized by input control. A-C, Seven-day-old MS grown Col-0 and *hac1-2* seedlings were subjected to 3h of HS treatment at 37°C. Data shown are representative of one biological replicates. Experiments were repeated twice with similar results.

**Supplemental Figure S10. Transcriptome comparison of TOR microarray with differentially expressed genes in***hac1* **and***gcn5* **mutants. A**, Venn diagram showing overlap of genes differentially expressed in *tor* and *hac1/5* mutants. Transcriptome data of TOR and HAC1 was obtained from public resources (Xiong *et al.* 2013; Jin *et al.* 2018). Genes in *hac1/5* mutants were extracted based on their false discovery rate value less than 0.05. **B, C**, GO biological process and molecular function of 269 genes commonly down-regulated by *tor* and *hac1/5*. **D**, Venn diagram showing overlap of co-expression genes between TOR, HAC1 and RAPTOR1B. Co-expression genes were extracted from ATTED-II database and total 2000 genes were used for co-expression genes overlap. **E**, GO biological process of 909 commonly co-expressed genes between TOR, HAC1 and RAPTOR1B. **F**, GO molecular function of 909 commonly co-expressed genes between TOR, HAC1 and RAPTOR1B. Panther 15.0 tool was used to analyse GO fold enrichment using Bonferroni correction and Fisher’s exact test type.

**Supplemental Figure S11. Protein-Protein interaction (PPI) between HAC1 and TOR complexes.** String data showing evidence of co-expression and interaction between TOR, RAPTOR1 and HAC1. Protein-Protein interaction (PPI) between HAC1 and TOR complexes was explored in String v.11 (Szklarczyk et al., 2019) which uses evidence from homologs in other species.

**Supplemental Figure S12. TOR remembers past exposure of heat stress. A**, Phenotype of Arabidopsis Col-0, G166, G548 and *tor35-7* after LAT HS. **B**, Percentage seedling survival and Chl estimation of Arabidopsis Col-0, G166, G548 and *tor35-7* after LAT HS. Five-day-old Arabidopsis seedlings were transferred to without or with Glc containing MS medium for 24h followed by HS. HS was applied as 1h_37ºC, 2d_22ºC, 2.5h_45ºC and 3-5d_22ºC. Data shown are representative of three biological replicates. Experiment were repeated thrice with similar results. Error bars represent SD (Student’s t test, P < 0.05; *, Control vs treatment; **, WT vs mutant/overexpression).

**Supplemental Figure S13. TOR overexpression lines primed with HS display enhanced thermotolerance. A**, Seedling survival phenotype of Col-0, G166 and G548 seedlings treated without or with Glc and subjected to unpriming and priming HS. Priming HS was applied as 1h_37ºC, 2h_22ºC, 2.5h_45ºC, 3-5d_22ºC. Unpriming HS was applied as 2.5h_45ºC, 3-5d_22ºC. **B**, Percentage seedling survival, fresh weight and Chl content in Arabidopsis Col-0, G166 and G548 seedlings after priming and unpriming HS. Five-day-old Arabidopsis Col-0, G166 and G548 seedlings were transferred to MS medium containing Glc (167mM) followed by unpriming and priming HS. Data shown are representative of three biological replicates. Experiments were repeated three times with similar results. Error bars represent SD (Student’s t test, P < 0.05; *, Control vs treatment; **, WT vs overexpression).

**Supplemental Figure S14. HS recovery under Glc induces***pTOR::GUS* **activity in the zones of primary meristems.** Picture showing *pTOR::GUS* activity after Glc and HS treatment. Seven-day-old MS grown Arabidopsis *pTOR::GUS* seedlings were transferred to without or with Glc (167mM) containing MS medium for 24h followed by HS. HS was applied as 1h_37ºC, 2h_22ºC, 2.5h_45ºC. Following HS, seedlings were recovered for 0h and 24h at 22ºC and transferred to GUS buffer. Images shown are representative of three biological replicates each containing more than four seedlings. Experiment were repeated thrice with similar results.

**Supplemental Figure S15. Arabidopsis***e2fa-1* **plants could not sustain thermomemory gene expression. A**, RT-qPCR expression of thermomemory related genes in Arabidopsis Col-0 and *e2fa-1* seedlings. Seven-day-old MS grown Arabidopsis Col-0 and *e2fa-1* seedlings containing Glc (56mM; under long-day conditions; 16h light and 8h dark, 100 μM m-2 s-1 light intensity) treated without or with HS. HS was applied as 1h_37ºC, 90min_22ºC, 45min_45ºC. After HS, seedlings were recovered for various time points (3h, 24h and 48h) in the culture room at 22ºC. Data shown are average of four technical replicates among two biological replicates. Experiments were repeated twice with similar results. Error bars represent SD (Student’s t test, P < 0.05; **, WT vs mutant).

**Supplemental Figure S16.** *e2fa-1* **mutants cause weaker LAT. A**, Chlorophyll estimation of Col-0, and *e2fa-1* seedlings after LAT. Five-day-old MS grown Col-0 and *e2fa-1* seedlings were transferred to without or with glucose containing MS medium for 24h followed by HS. HS was applied as 1h_37ºC, 2d_22ºC, 2.5h_45ºC and 3-5_22ºC. Data shown are average of three biological replicates. Experiments were repeated thrice with similar results. **B**, Images showing HS phenotype of Col-0 and *e2fa-1* after LAT. **(C)** Images showing HS phenotype of Col-0 and e2fa-1 after LAT. **C**, ChIP-qPCR showing binding of E2Fa at thermomemory gene promoters. Seven-day-old Arabidopsis Col-0 plants were used. Amount of immuno-precipitated promoter DNA was calculated by comparing samples treated without or with anti-E2Fa serum. CT Values without and with anti-E2Fa serum were normalized by input control. MCM5 and ATXR6 are known E2Fa target gene and were taken as positive control. Data shown are representative of four technical replicates among two biological replicates. Experiments were repeated twice with similar results. Error bars represent SD (Student’s t test, P < 0.05; *, Control vs treatment; **, WT vs mutant).

**Supplemental Figure S17. Putative TOR-E2Fa interactions with chromatin machinery. A** Network showing protein interaction of E2Fa with BRM, SYD and SWI3C. Data was extracted from BIOGRID (Efroni *et al.* 2013). **B**, Table showing sucrose/TOR phosphorylation of proteins involved in chromatin remodelling/histone synthesis and modification. Data was extracted from a list of proteins regulated by sucrose/TOR (Van Leene *et al.* 2019). Colour showing peptide numbers obtained from the given list of proteins. High red intensity showing maximum number of peptides for a given protein whereas green colour is showing least number. Star denotes as significance level representing more than two peptides in each protein.

**Supplemental Figure S18. Similar to TOR, RAPTOR1B shares huge overlap with BRM. A**, GO biological process of 966 common genes between TOR, BRM and RAPTOR1B. **B**, GO molecular function of 966 common genes between TOR, BRM and RAPTOR1B. **C**, Venn diagram showing overlap of co-expression genes between TOR, BRM and RAPTOR1B. **D**, Venn diagram showing overlap of co-expression genes between TOR, BRM and E2Fa. Co-expression genes were extracted from ATTED-II database and total 2000 genes were used for co-expression genes overlap. **E**, GO biological process of 173 commonly co-expressed genes between TOR, BRM and E2Fa. **F**, GO molecular function of 173 commonly co-expressed genes between TOR, BRM and E2Fa. Panther 15.0 tool was used to analyse GO fold enrichment using Bonferroni correction and Fisher’s exact test type.

**Supplemental Figure S19. Arabidopsis TOR co-expression shares huge overlap with chromatin remodeler BRAHMA and SYD (CHR3). A**, GO biological process enrichment of 1236 genes commonly co-expressed between TOR, BRM and SYD. **B**, GO molecular function enrichment of 1236 genes commonly co-expressed between TOR, BRM and SYD. **C**, Venn diagram showing overlap of BRM and REF6 ChIP-seq targets with genes containing E2F binding motifs in their promoters. **D**, GO biological process enrichment of 1284 genes between BRM targets and E2F motifs containing genes. **E**, GO molecular function enrichment of 1284 genes between BRM targets and E2F motifs containing genes. BRM and REF6 ChIP-seq data was obtained from publicly available resources (Li *et al.* 2016). Panther 15.0 tool was used to analyse GO fold enrichment using Bonferroni correction and Fisher’s exact test type.

