## Supplemental Figures for "TOR coordinates with transcriptional and chromatin machinery to regulate thermotolerance and thermomemory"

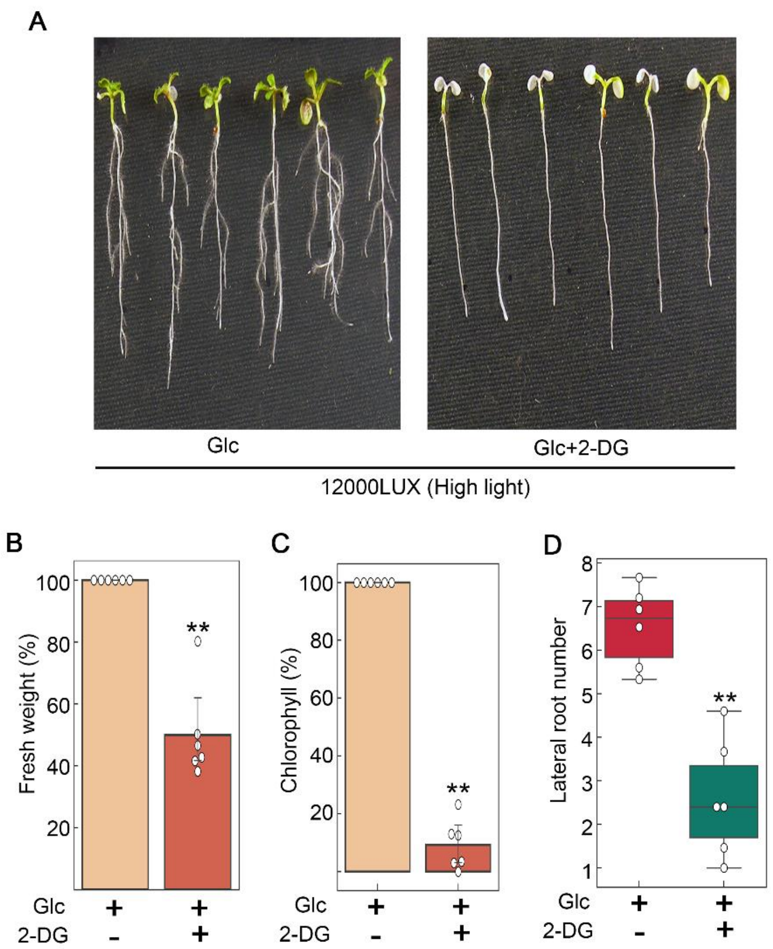


**Supplemental Figure S1. Photosynthesis generated Glc could not provide thermotolerance under high light intensity when blocked with Glc analogue 2-deoxy-glucose. A,** Phenotype showing effect of Glc analogue 2-DG on thermotolerance response. **B-D,** Percentage fresh weight, chlorophyll content and lateral root numbers of Arabidopsis Col-0 seedlings after treatment with high light and heat stress in presence of Glc alone and Glc along with its analogue 2-DG. Five-day-old MS grown Arabidopsis Col-0 seedlings under standard light intensity were transferred to Glc (56mM) containing MS media without or with Glc analogue 2-DG (2.5mM). After transfer, seedlings were transferred to high light intensity (approximately 12000 LUX) for 24h and then subjected to heat stress at 1h_37ºC, 2h_22ºC, 2.5h_45ºC and 3-7d_22ºC. Data shown are average of three biological replicates. Experiment was repeated thrice with similar results. Error bars represent SD (Student's t test, $P < 0.05$; **, Mock vs inhibitor).

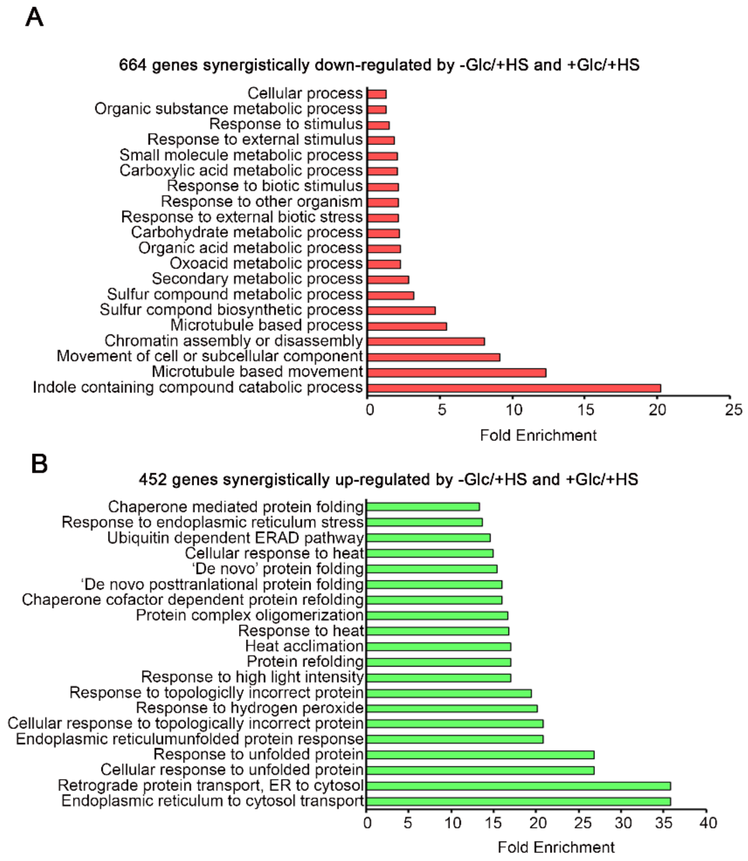


**Supplemental Figure S2. Gene ontology of common genes between –Glc/+HS and +Glc/+HS exhibits up-regulation of genes involved in protein folding. A, B** GO term enrichment of biological process synergistically down-and up-regulated by both –Glc/+HS and +Glc/+HS. Top twenty GO categories were included based on their fold enrichment. Panther 15.0 tool was used to analyse GO fold enrichment using Bonferroni correction and Fisher's exact test type.

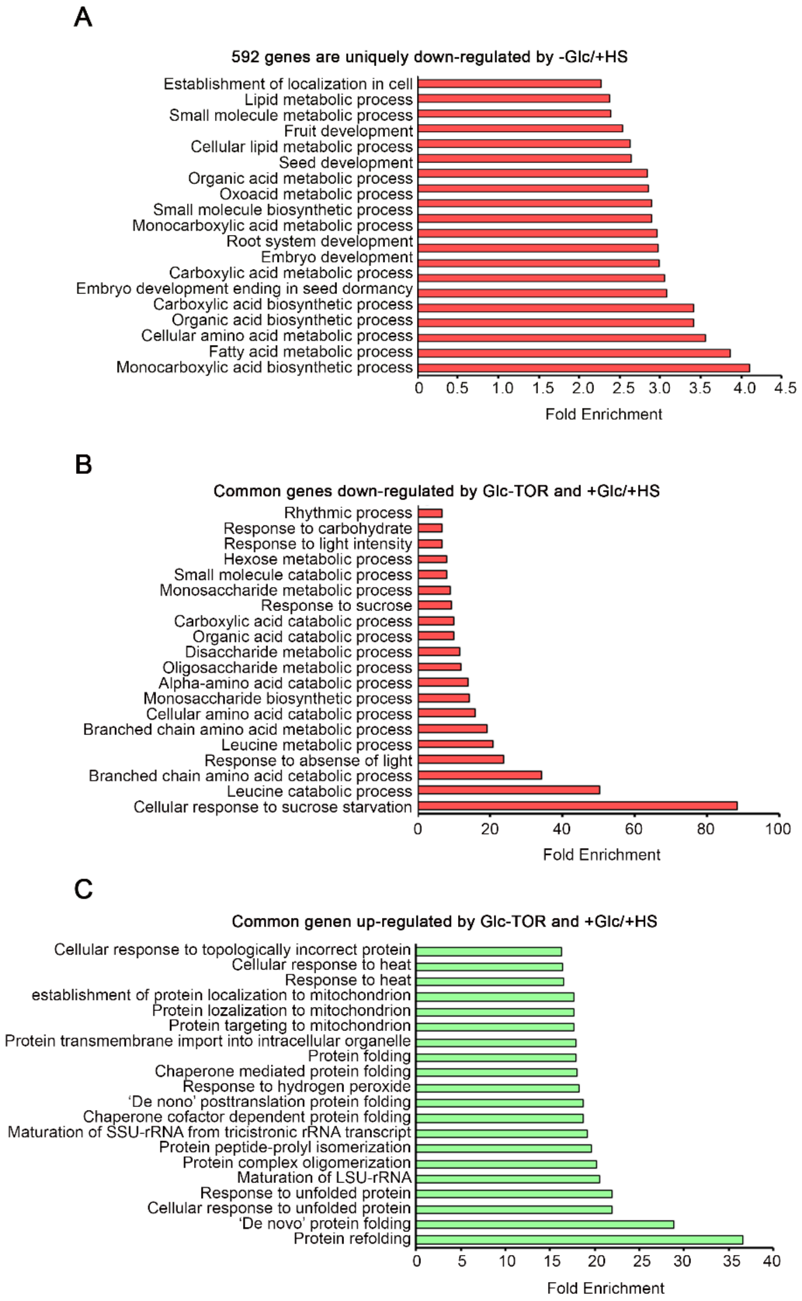


**Supplemental Figure S3. Glc regulates HS transcriptome largely through TOR pathway. A,** GO biological process of genes exclusively down-regulated by –Glc/+HS. **B, C,** GO biological process of genes commonly down- and up-regulated by +Glc/+HS and public available Glc-TOR microarray genes (Xiong *et al.* 2013). Top twenty GO categories were included based on their fold enrichment. Panther 15.0 tool was used to analyse GO fold enrichment using Bonferroni correction and Fisher's exact test type.

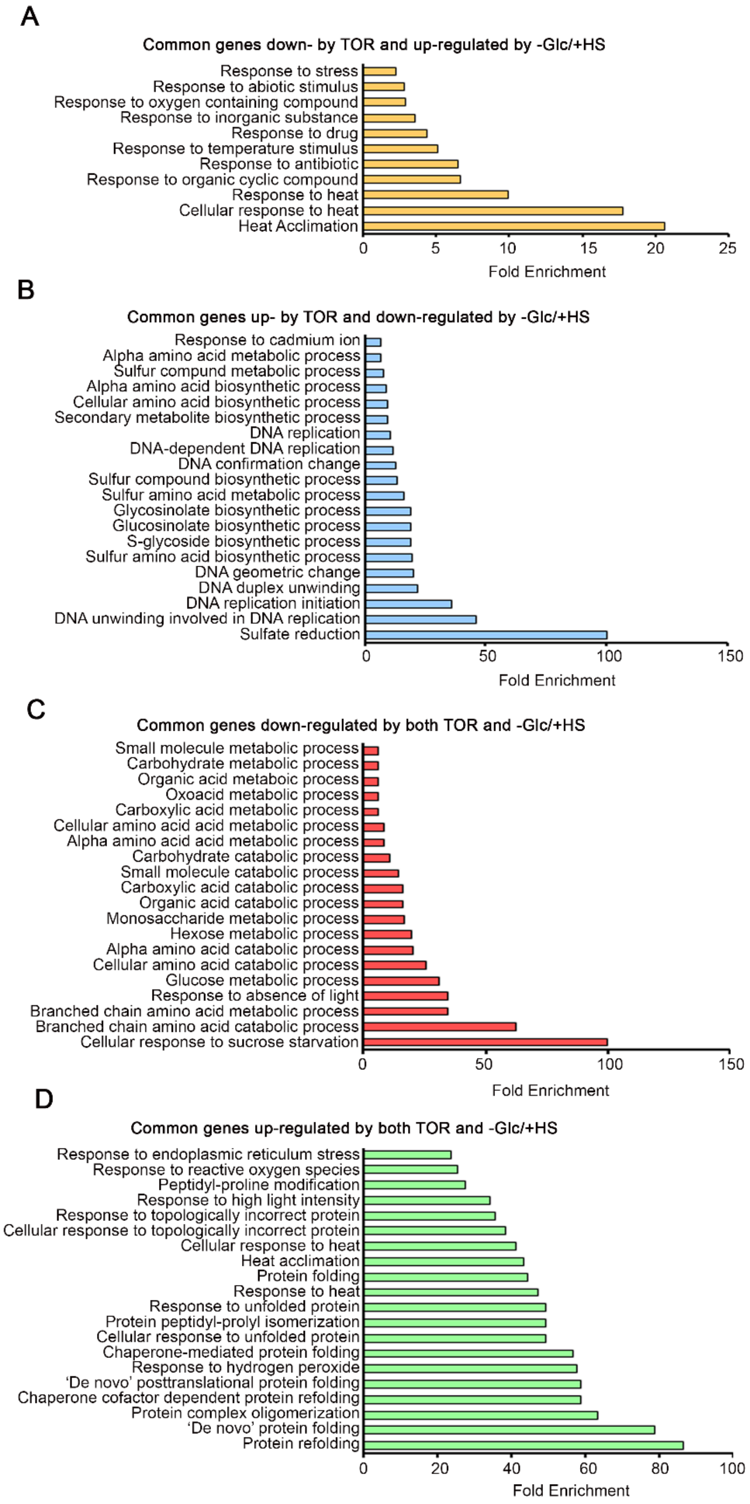


**Supplemental Figure S4. HS transcriptome induced in absence of Glc exhibits large antagonistic interaction with Glc-TOR target genes. A, B,** GO biological process of genes antagonistically (down- by Glc-TOR and up-regulated by –Glc/+HS; up- by Glc-TOR and down-regulated by –Glc/+HS) regulated by Glc-TOR and –Glc/+HS. **C, D,** GO biological process enrichment of genes synergistically (up- and down-regulated by both Glc-TOR and –Glc/+HS) regulated by Glc-TOR and –Glc/+HS. Top twenty GO categories were included based on their fold enrichment. Panther 15.0 tool was used to analyse GO fold enrichment using Bonferroni correction and Fisher's exact test type.

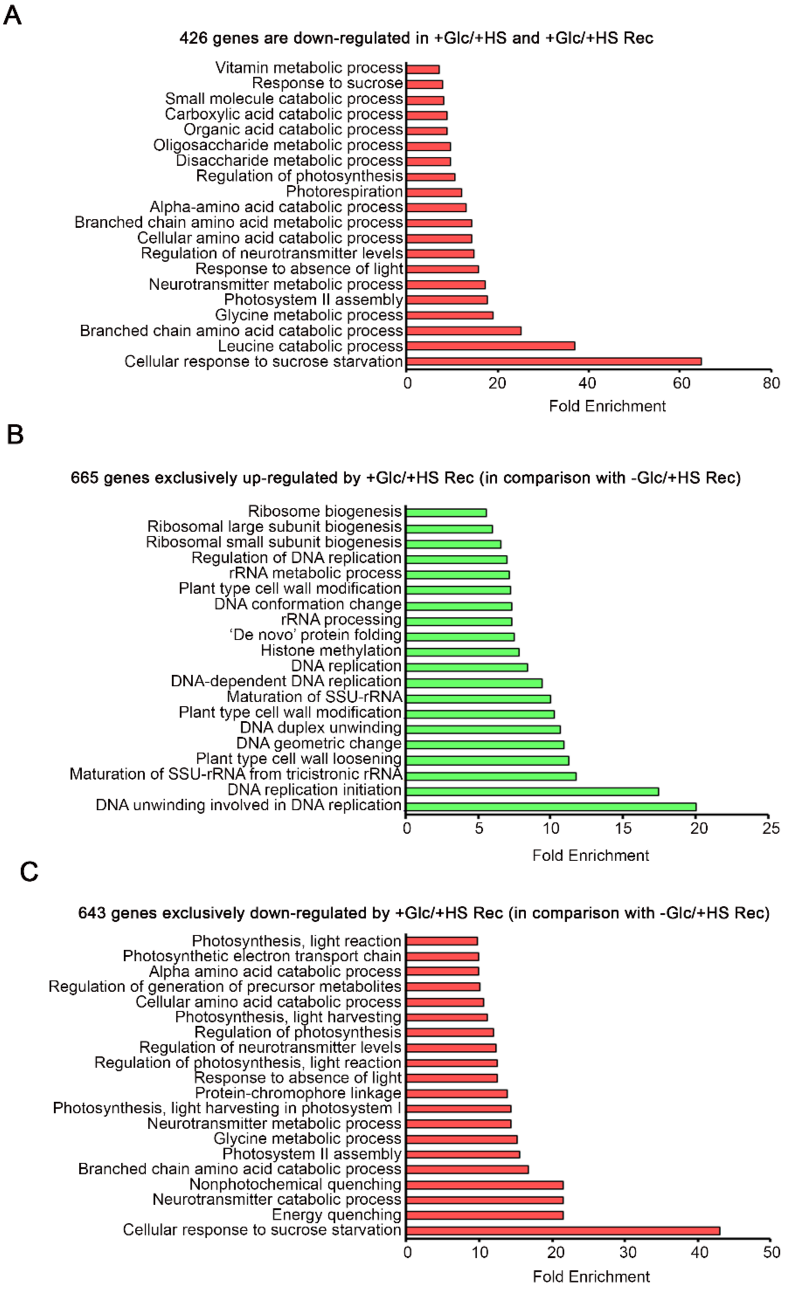


**Supplemental Figure S5. Presence of Glc regulates sustenance of HS transcriptome for HS recovery. A,** GO biological process of genes commonly down-regulated by +Glc/+HS and +Glc/+HS Rec. **B, C,** GO biological process of genes exclusively up- and down-regulated by +Glc/+HS Rec in comparison with –Glc/+HS Rec. Top twenty GO categories were included based on their fold enrichment. Panther 15.0 tool was used to analyse GO fold enrichment using Bonferroni correction and Fisher's exact test type.

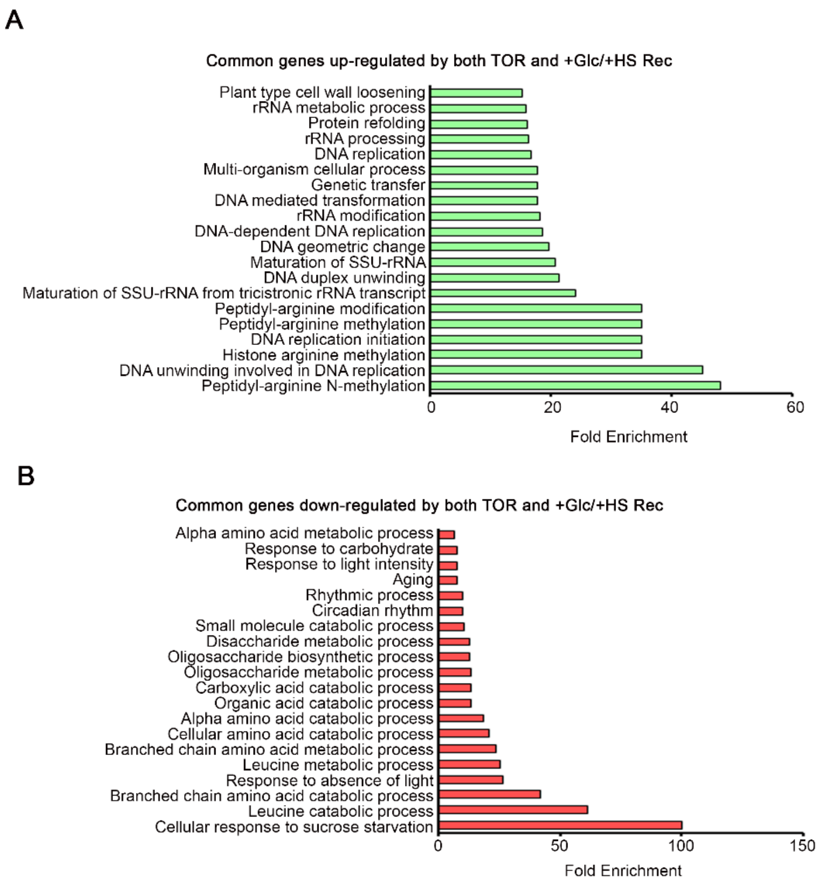


**Supplemental Figure S6. HS recovery under Glc induces processes required for growth/HS recovery in a TOR dependent manner. A,** GO biological process of genes commonly up-regulated by +Glc/+HS Rec and TOR. **B,** GO biological process of genes commonly down-regulated by +Glc/+HS Rec and TOR. Top twenty GO categories were included based on their fold enrichment. Panther 15.0 tool was used to analyse GO fold enrichment using Bonferroni correction and Fisher's exact test type.

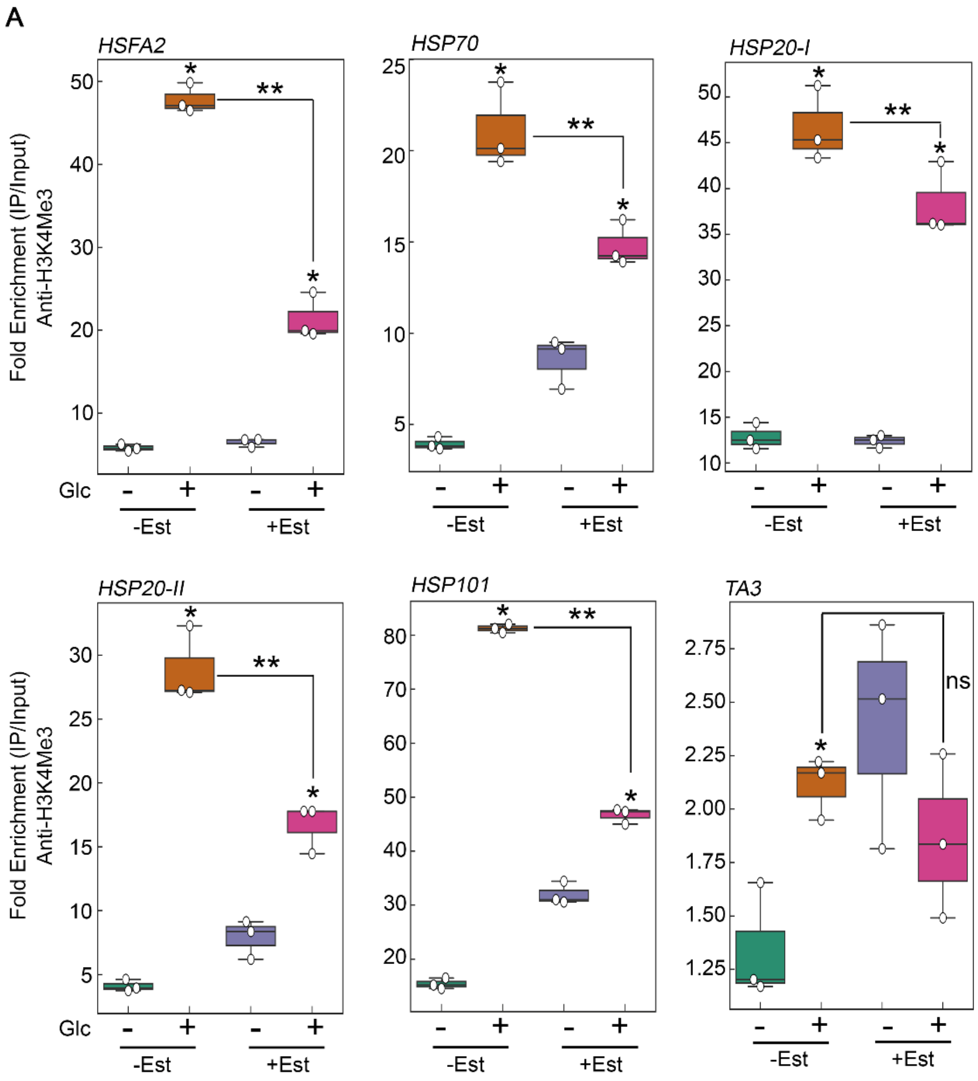


**Supplemental Figure S7. Glc through TOR induces enrichment of histone H3K4me2 marks at the promoters of HS genes.** ChIP-qPCR showing enrichment of histone H3K4me2 at the promoters of HS loci under Glc lacking/sufficiency. Promoter fragments containing cis-acting HSEs were immuno-precipitated using anti-H3K4me2 antibody. Amount of immuno-precipitated promoter DNA was calculated by comparing samples treated without or with anti-H3K4me2 antibody. Ct values without and with antibody samples were normalized by input control. TA3 is a highly heterochromatinized DNA and was used as a negative control. Five-day-old *tor-es1* seedlings were transferred to MS medium containing 20µM β-estradiol for four days. Following estradiol treatment, seedlings were subjected to 24h energy starvation in MS medium without Glc and then supplied with 3h Glc treatment. For mock treatment, seedlings were transferred in equal volume of DMSO (as used for estradiol) containing MS medium. Data shown are representative of one biological replicates. Experiments were repeated twice with similar results. Error bars = SD (Student's t test, P, 0.05; *control versus treatment; **Mock versus Estradiol).

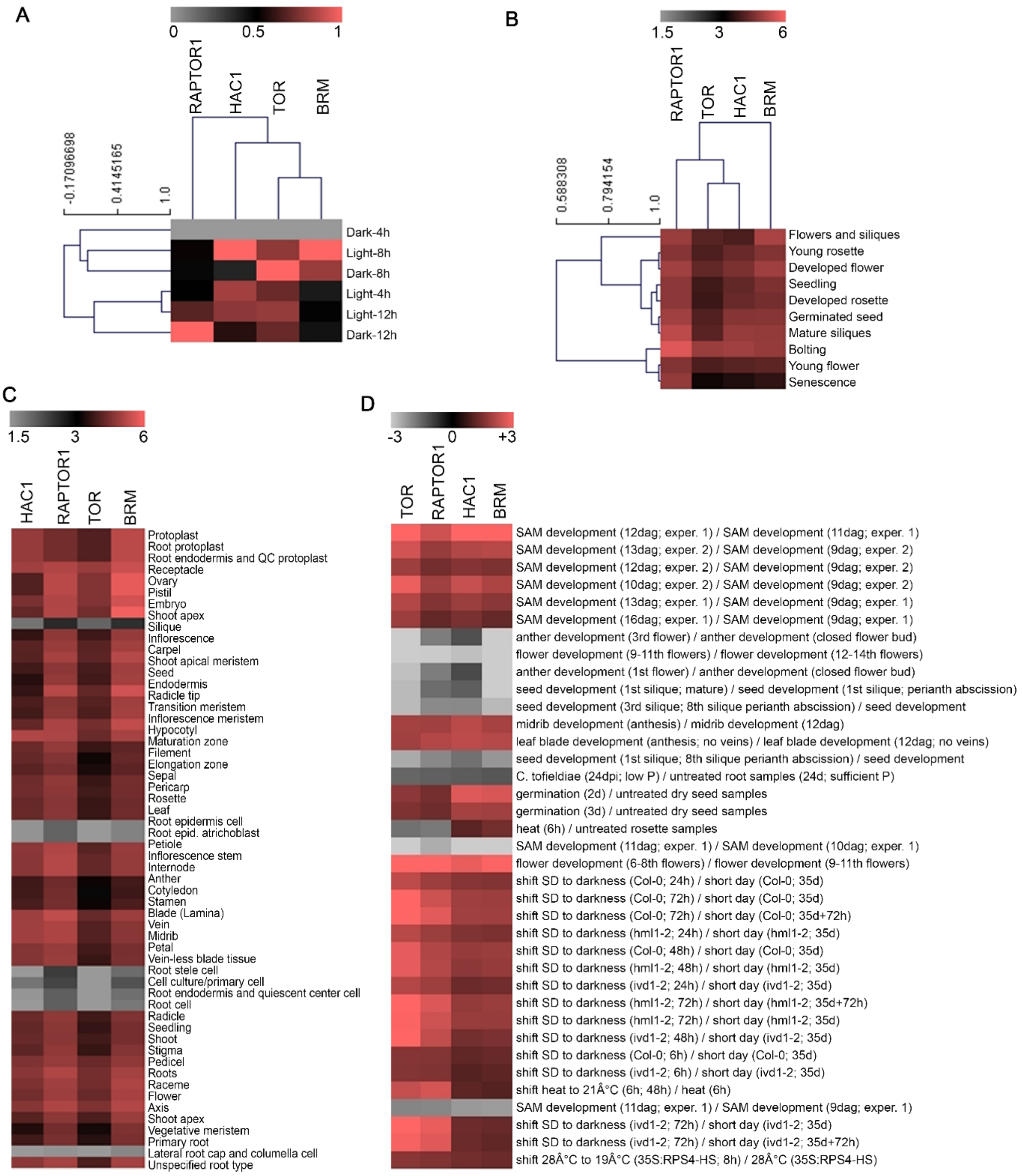


**Supplemental Figure S8. Diurnal light-dark cycle and Genevestigator transcriptome comparison shows similarity in expression between TOR, RAPTOR1, HAC1 and BRM. A,** Heat map showing expression profiling of TOR, RAPTOR1, HAC1 and BRM in diurnal light-dark transcriptome data obtained from public resources (Ferrari *et al.* 2019). **B-D,** Heat maps showing expression profiling of TOR, RAPTOR1, HAC1 and BRM in anatomy, development and perturbations data obtained from Genevestigator.

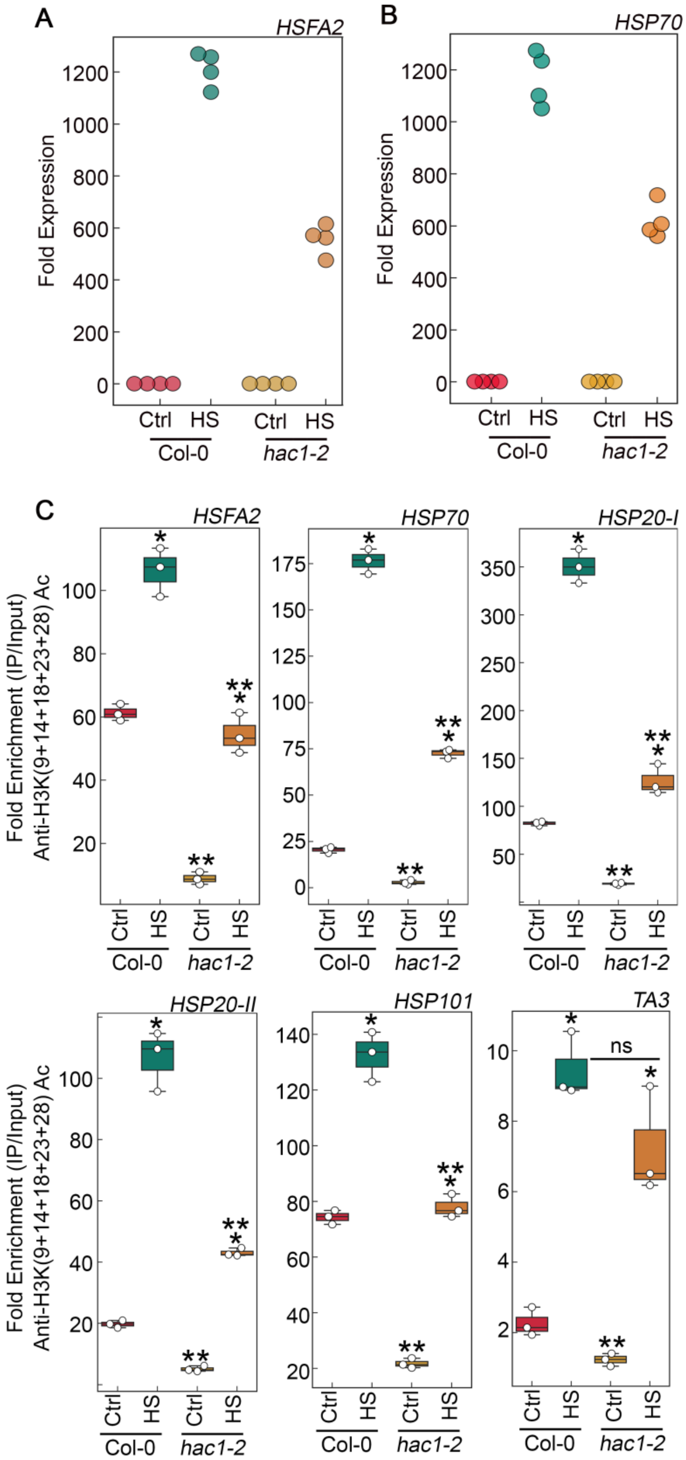


**Supplemental Figure S9. *hac1* mutant demonstrates less accumulation of H3K acetylation, leading to perturbation in HS induced gene expression. A, B,** RT-qPCR expression of HS genes in Col-0 and *hac1* mutants in response to HS. **C,** ChIP-qPCR showing enrichment of histone H3K Ac (9+14+18+ 23+27) at the promoters of HS gene encompassing HSEs in Col-0 and *hac1-2* mutant. Amount of immuno-precipitated promoter DNA was calculated by comparing samples treated without or with anti-Histone H3K (9+14+18+ 23+27) acetyl antibody. Ct values without and with antibody samples were normalized by input control. A-C, Seven-day-old MS grown Col-0 and *hac1-2* seedlings were subjected to 3h of HS treatment at 37°C. Data shown are representative of one biological replicates. Experiments were repeated twice with similar results.

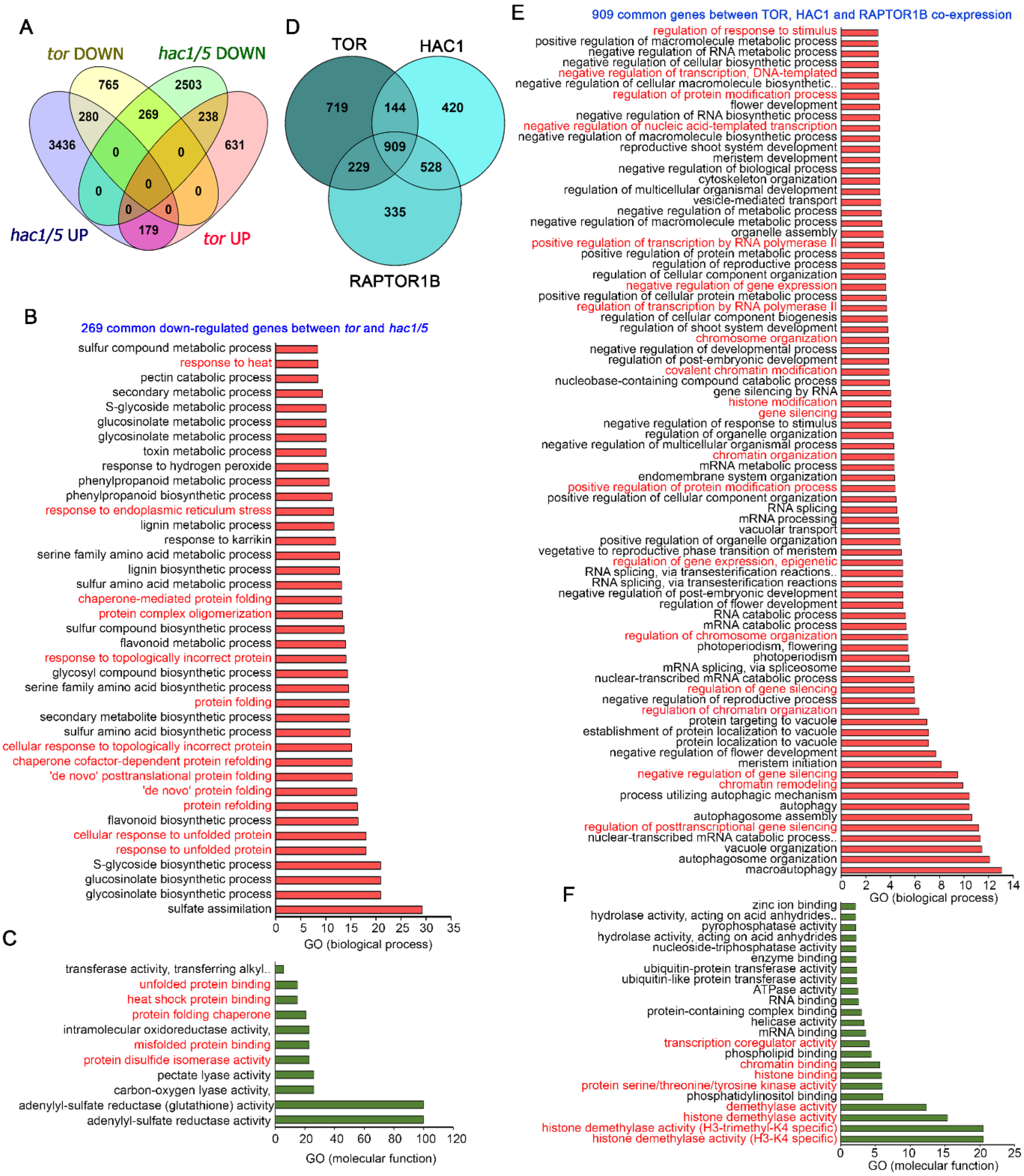

**Supplemental Figure S10. Transcriptome comparison of TOR microarray with differentially expressed genes in *hac1* and *gcn5* mutants. A,** Venn diagram showing overlap of genes differentially expressed in *tor* and *hac1/5* mutants. Transcriptome data of TOR and HAC1 was obtained from public resources (Xiong *et al.* 2013; Jin *et al.* 2018). Genes in *hac1/5* mutants were extracted based on their false discovery rate value less than 0.05. **B, C,** GO biological process and molecular function of 269 genes commonly down-regulated by *tor* and *hac1/5*. **D,** Venn diagram showing overlap of co-expression genes between TOR, HAC1 and RAPTOR1B. Co-expression genes were extracted from ATTED-II database and total 2000 genes were used for co-expression genes overlap. **E,** GO biological process of 909 commonly co-expressed genes between TOR, HAC1 and RAPTOR1B. **F,** GO molecular function of 909 commonly co-expressed genes between TOR, HAC1 and RAPTOR1B. Panther 15.0 tool was used to analyse GO fold enrichment using Bonferroni correction and Fisher's exact test type.

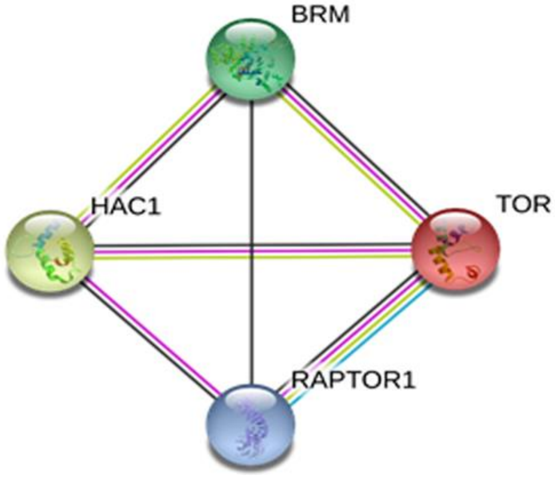


**Supplemental Figure S11. Protein-Protein interaction (PPI) between HAC1 and TOR complexes.** String data showing evidence of co-expression and interaction between TOR, RAPTOR1 and HAC1. Protein-Protein interaction (PPI) between HAC1 and TOR complexes was explored in String v.11 (Szklarczyk et al., 2019) which uses evidence from homologs in other species.

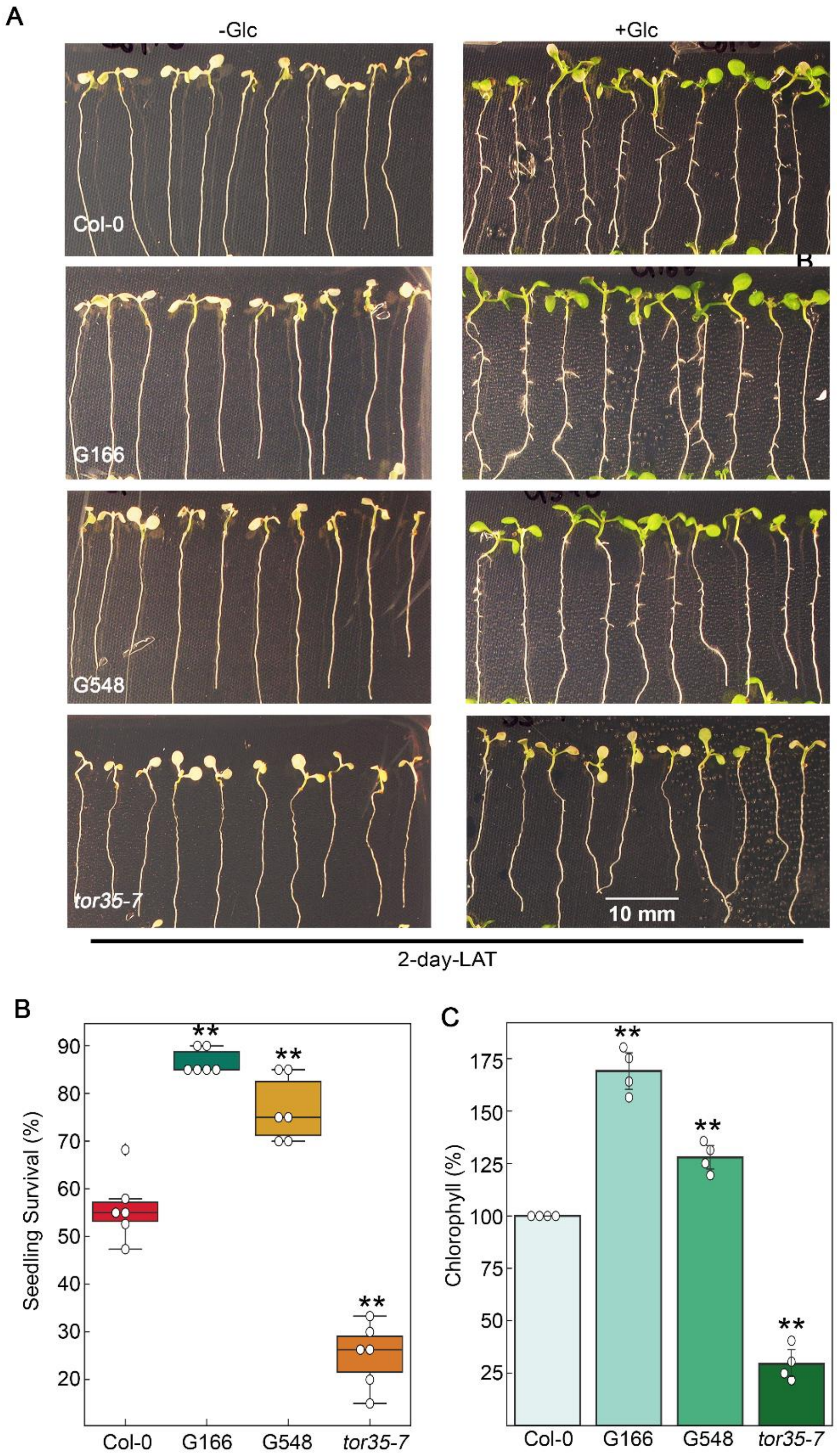


**Supplemental Figure S12. TOR remembers past exposure of heat stress. A,** Phenotype of Arabidopsis Col-0, G166, G548 and *tor35-7* after LAT HS. **B,** Percentage seedling survival and Chl estimation of Arabidopsis Col-0, G166, G548 and *tor35-7* after LAT HS. Five-day-old Arabidopsis seedlings were transferred to without or with Glc containing MS medium for 24h followed by HS. HS was applied as 1h_37ºC, 2d_22ºC, 2.5h_45ºC and 3-5d_22ºC. Data shown are representative of three biological replicates. Experiment were repeated thrice with similar results. Error bars represent SD (Student's t test, $P < 0.05$; *, Control vs treatment; **, WT vs mutant/overexpression).

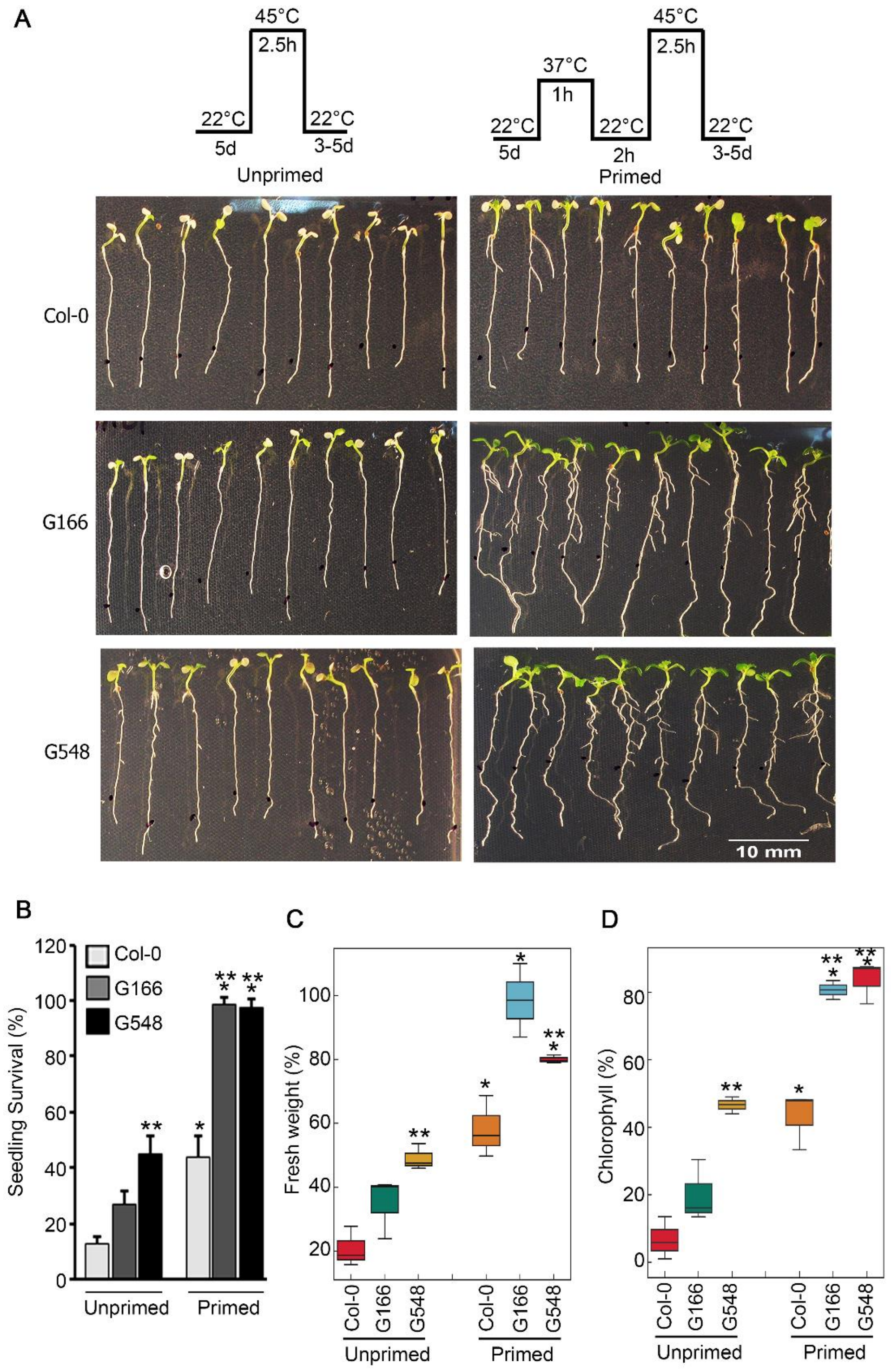


**Supplemental Figure S13. TOR overexpression lines primed with HS display enhanced thermotolerance. A,** Seedling survival phenotype of Col-0, G166 and G548 seedlings treated without or with Glc and subjected to unpriming and priming HS. Priming HS was applied as 1h_37ºC, 2h_22ºC, 2.5h_45ºC, 3-5d_22ºC. Unpriming HS was applied as 2.5h_45ºC, 3-5d_22ºC. **B,** Percentage seedling survival, fresh weight and Chl content in Arabidopsis Col-0, G166 and G548 seedlings after priming and unpriming HS. Five-day-old Arabidopsis Col-0, G166 and G548 seedlings were transferred to MS medium containing Glc (167mM) followed by unpriming and priming HS. Data shown are representative of three biological replicates.

Experiments were repeated three times with similar results. Error bars represent SD (Student's t test, P < 0.05; *, Control vs treatment; **, WT vs overexpression).

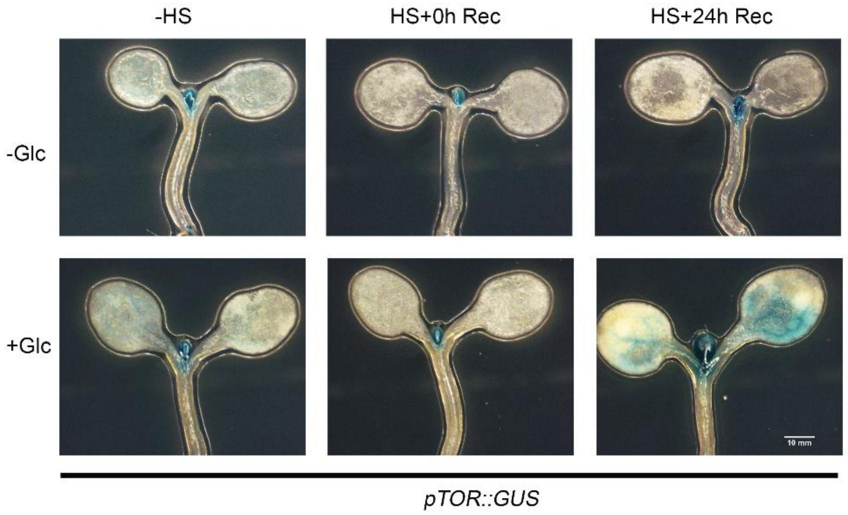


**Supplemental Figure S14. HS recovery under Glc induces *pTOR::GUS* activity in the zones of primary meristems.** Picture showing *pTOR::GUS* activity after Glc and HS treatment. Seven-day-old MS grown Arabidopsis *pTOR::GUS* seedlings were transferred to without or with Glc (167mM) containing MS medium for 24h followed by HS. HS was applied as 1h_37ºC, 2h_22ºC, 2.5h_45ºC. Following HS, seedlings were recovered for 0h and 24h at 22ºC and transferred to GUS buffer. Images shown are representative of three biological replicates each containing more than four seedlings. Experiment were repeated thrice with similar results.

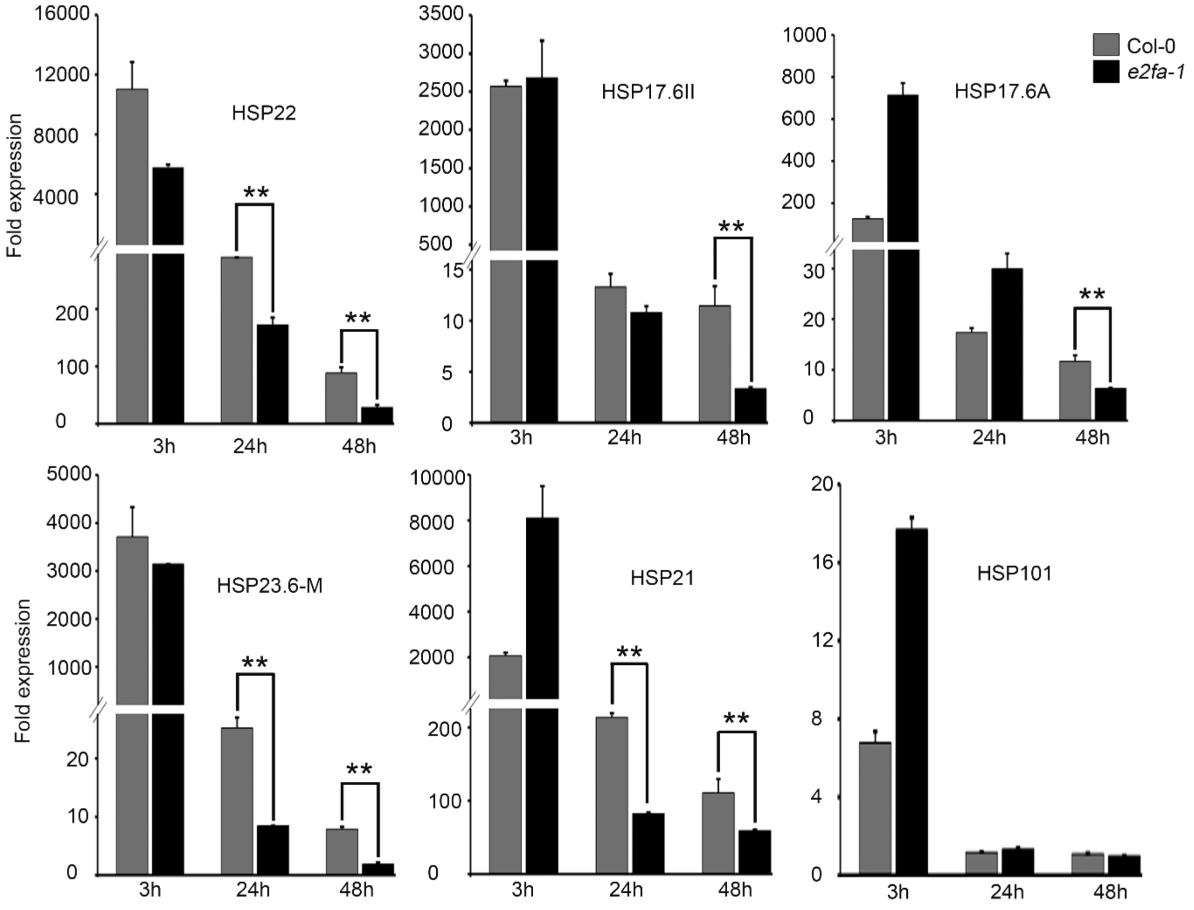


**Supplemental Figure S15. Arabidopsis *e2fa-1* plants could not sustain thermomemory gene expression. A,** RT-qPCR expression of thermomemory related genes in Arabidopsis Col-0 and *e2fa-1* seedlings. Seven-day-old MS grown Arabidopsis Col-0 and *e2fa-1* seedlings containing Glc (56mM; under long-day conditions; 16h light and 8h dark, 100 µM m-2 s-1 light intensity) treated without or with HS. HS was applied as 1h_37ºC, 90min_22ºC, 45min_45ºC. After HS, seedlings were recovered for various time points (3h, 24h and 48h) in the culture room at 22ºC. Data shown are average of four technical replicates among two biological replicates. Experiments were repeated twice with similar results. Error bars represent SD (Student's t test, $P < 0.05$; **, WT vs mutant).

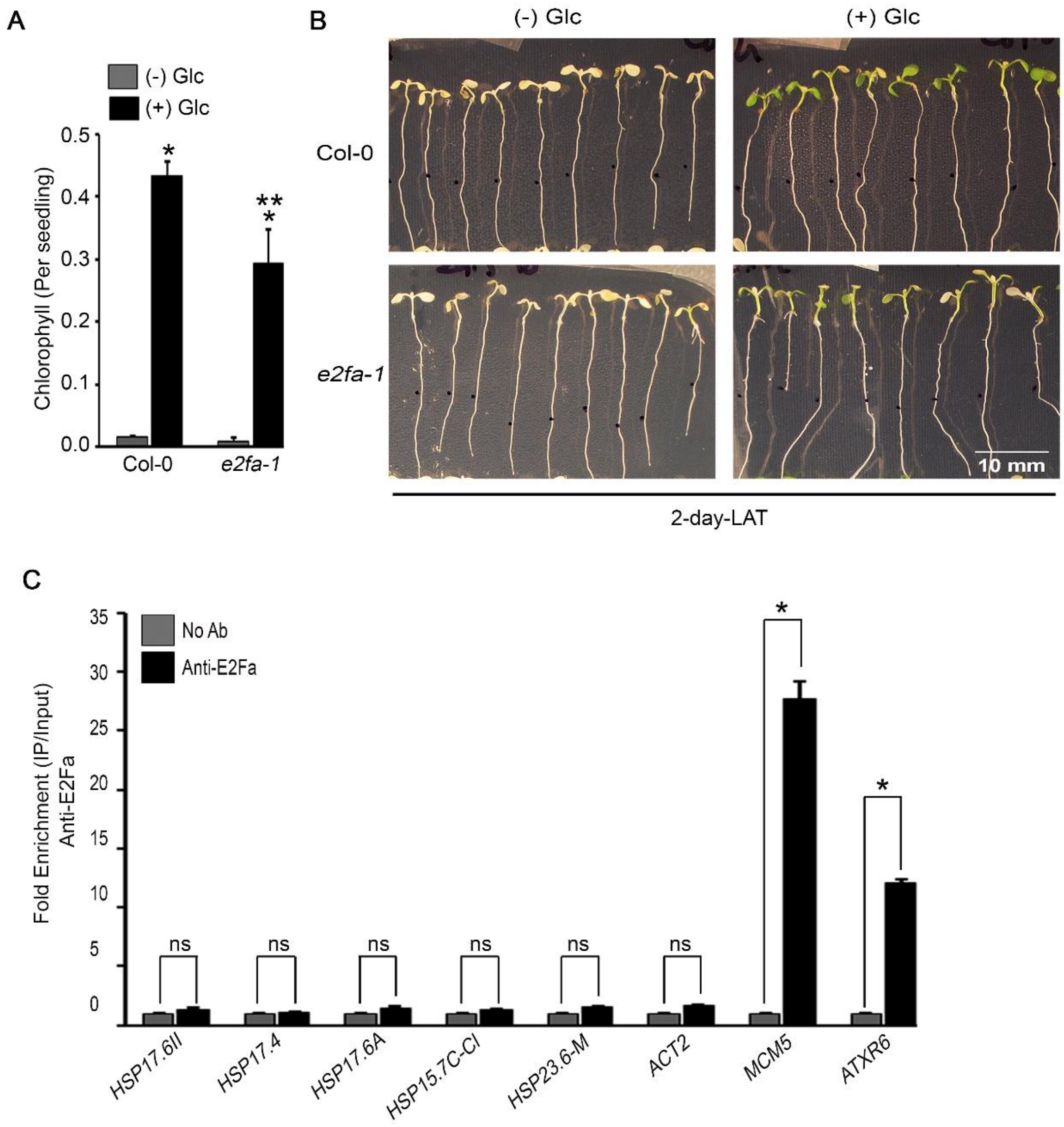


**Supplemental Figure S16. *e2fa-1* mutants cause weaker LAT. A,** Chlorophyll estimation of Col-0, and *e2fa-1* seedlings after LAT. Five-day-old MS grown Col-0 and *e2fa-1* seedlings were transferred to without or with glucose containing MS medium for 24h followed by HS. HS was applied as 1h_37ºC, 2d_22ºC, 2.5h_45ºC and 3-5_22ºC. Data shown are average of three biological replicates. Experiments were repeated thrice with similar results. **B,** Images showing HS phenotype of Col-0 and *e2fa-1* after LAT. **(C)** Images showing HS phenotype of Col-0 and e2fa-1 after LAT. **C,** ChIP-qPCR showing binding of E2Fa at thermomemory gene promoters. Seven-day-old Arabidopsis Col-0 plants were used. Amount of immuno-precipitated promoter DNA was calculated by comparing samples treated without or with anti-E2Fa serum. CT Values without and with anti-E2Fa serum were normalized by input control. MCM5 and ATXR6 are known E2Fa target gene and were taken as positive control. Data shown are representative of four technical replicates among two biological replicates. Experiments were repeated twice with similar results. Error bars represent SD (Student's t test, $P < 0.05$; *, Control vs treatment; **, WT vs mutant).

A

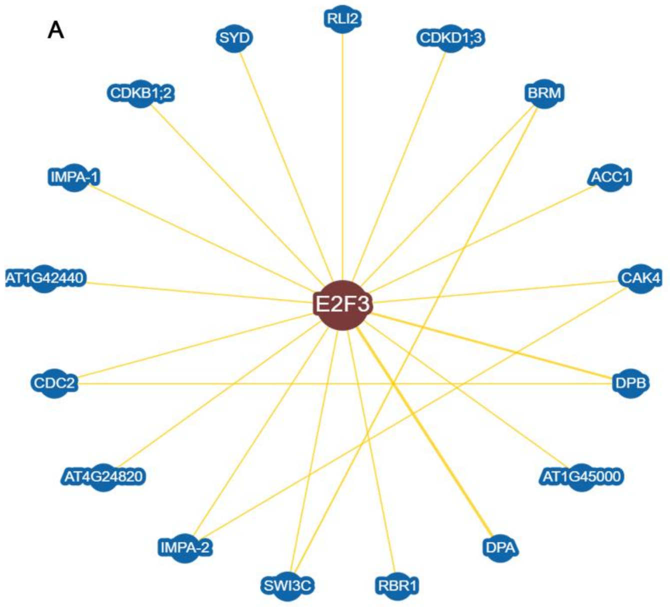


B

| 26 chromatin remodelers containing 83 phosphopeptides | | |
|---|---|---|
| **Gene ID** | **Gene Name** | **Number of peptides** |
| AT1G32750 | HAC13 protein (HAC13) | 2 |
| AT1G76710 | SET domain group 26;SET domain group 26 | 1 |
| AT1G77180 | chromatin protein family;chromatin protein family;chromatin protein family | 5* |
| AT1G77300 | histone methyltransferases(H3-K4 specific);histone methyltransferases(H3-K36 specific) | 2 |
| AT1G79000 | histone acetyltransferase of the CBP family 1;histone acetyltransferase of the CBP family 1 (HAC1) | 1 |
| AT2G13370 | chromatin remodeling 5 | 3* |
| AT2G16390 | SNF2 domain-containing protein / helicase domain-containing protein | 1 |
| AT2G25170 | chromatin remodeling factor CHD3 (PICKLE) | 2 |
| AT2G46020 | transcription regulatory protein SNF2, putative;transcription regulatory protein SNF2, putative, BRAHMA (BRM) | 15* |
| AT2G28290 | SWI2/SNF2-like protein in the SNF2 subclass, CHROMATIN REMODELING COMPLEX SUBUNIT R 3, CHR3 | 8* |
| AT3G12810 | SNF2 domain-containing protein / helicase domain-containing protein | 4* |
| AT3G19080 | SWIB complex BAF60b domain-containing protein | 3* |
| AT3G48430 | relative of early flowering 6 | 7* |
| AT3G54610 | histone acetyltransferase of the GNAT family 1 (GCN5) | 5* |
| AT4G15180 | SET domain protein 2 | 5* |
| AT4G29510 | arginine methyltransferase 11 | 1 |
| AT4G38130 | histone deacetylase 1;histone deacetylase 1 | 1 |
| AT5G18620 | chromatin remodeling factor17;chromatin remodeling factor17 | 1 |
| AT5G22220 | E2F transcription factor 1;E2F transcription factor 1 | 2 |
| AT5G22650 | histone deacetylase 2B;histone deacetylase 2B | 5* |
| AT5G27670 | histone H2A 7 | 2 |
| AT5G44800 | chromatin remodeling 4 | 1 |
| AT5G56740 | histone acetyltransferase of the GNAT family 2 | 1 |
| AT5G63950 | chromatin remodeling 24 | 2 |
| AT2G16440 | Minichromosome maintenance (MCM2/3/5) family protein | 2 |
| AT4G02060 | Minichromosome maintenance (MCM2/3/5) family protein;Minichromosome maintenance (MCM2/3/5) family protein | 1 |

**Supplemental Figure S17. Putative TOR-E2Fa interactions with chromatin machinery. A** Network showing protein interaction of E2Fa with BRM, SYD and SWI3C. Data was extracted from BIOGRID (Efroni *et al.* 2013). **B,** Table showing sucrose/TOR phosphorylation of proteins involved in chromatin remodelling/histone synthesis and modification. Data was extracted from a list of proteins regulated by sucrose/TOR (Van Leene *et al.* 2019). Colour showing peptide numbers obtained from the given list of proteins. High red intensity showing maximum number of peptides for a given protein whereas green colour is showing least number. Star denotes as significance level representing more than two peptides in each protein.

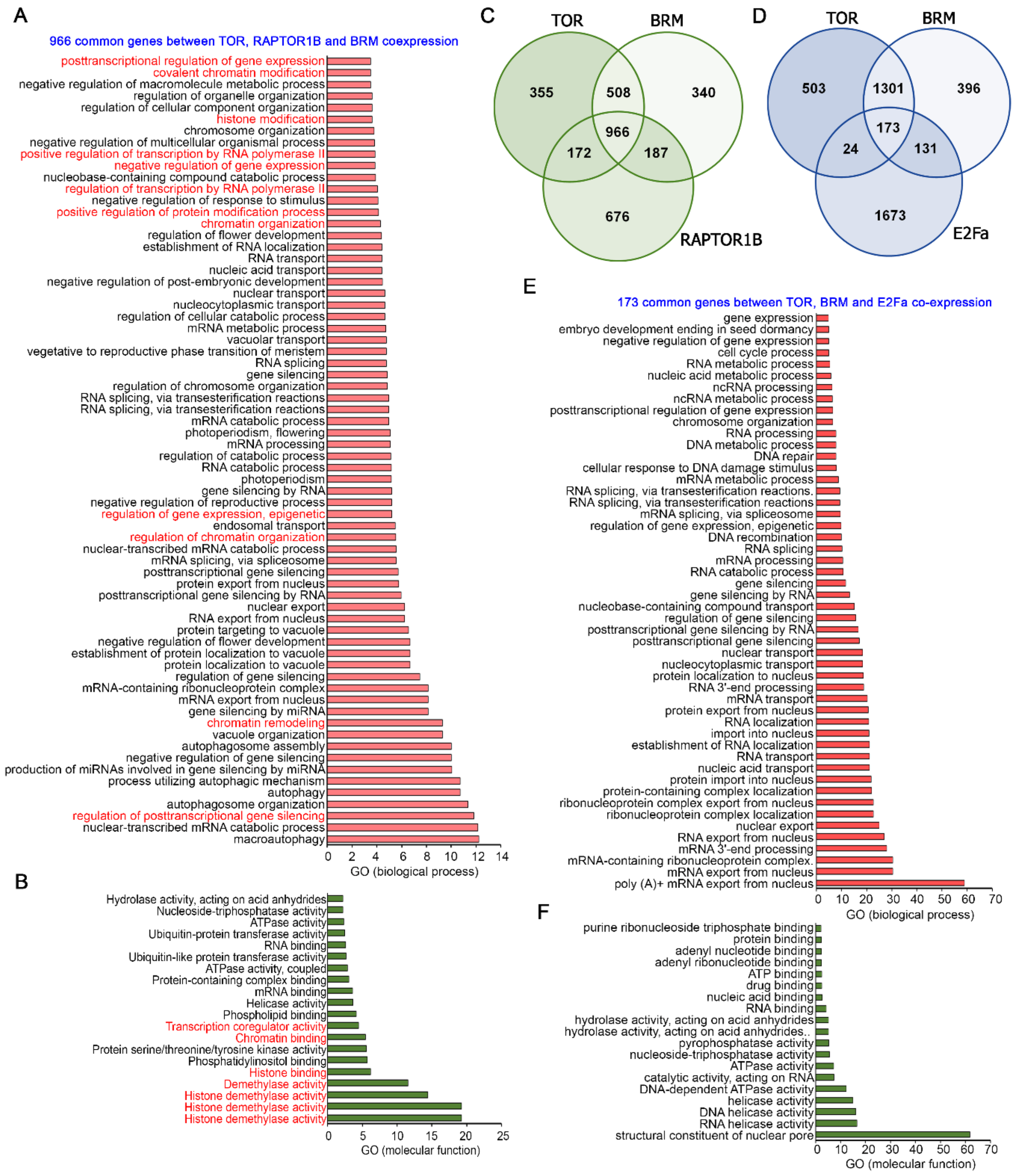


**Supplemental Figure S18. Similar to TOR, RAPTOR1B shares huge overlap with BRM. A,** GO biological process of 966 common genes between TOR, BRM and RAPTOR1B. **B,** GO molecular function of 966 common genes between TOR, BRM and RAPTOR1B. **C,** Venn diagram showing overlap of co-expression genes between TOR, BRM and RAPTOR1B. **D,** Venn diagram showing overlap of co-expression genes between TOR, BRM and E2Fa. Co-expression genes were extracted from ATTED-II database and total 2000 genes were used for co-expression genes overlap. **E,** GO biological process of 173 commonly co-expressed genes between TOR, BRM and E2Fa. **F,** GO molecular function of 173 commonly co-expressed genes between TOR, BRM and E2Fa. Panther 15.0 tool was used to analyse GO fold enrichment using Bonferroni correction and Fisher's exact test type.

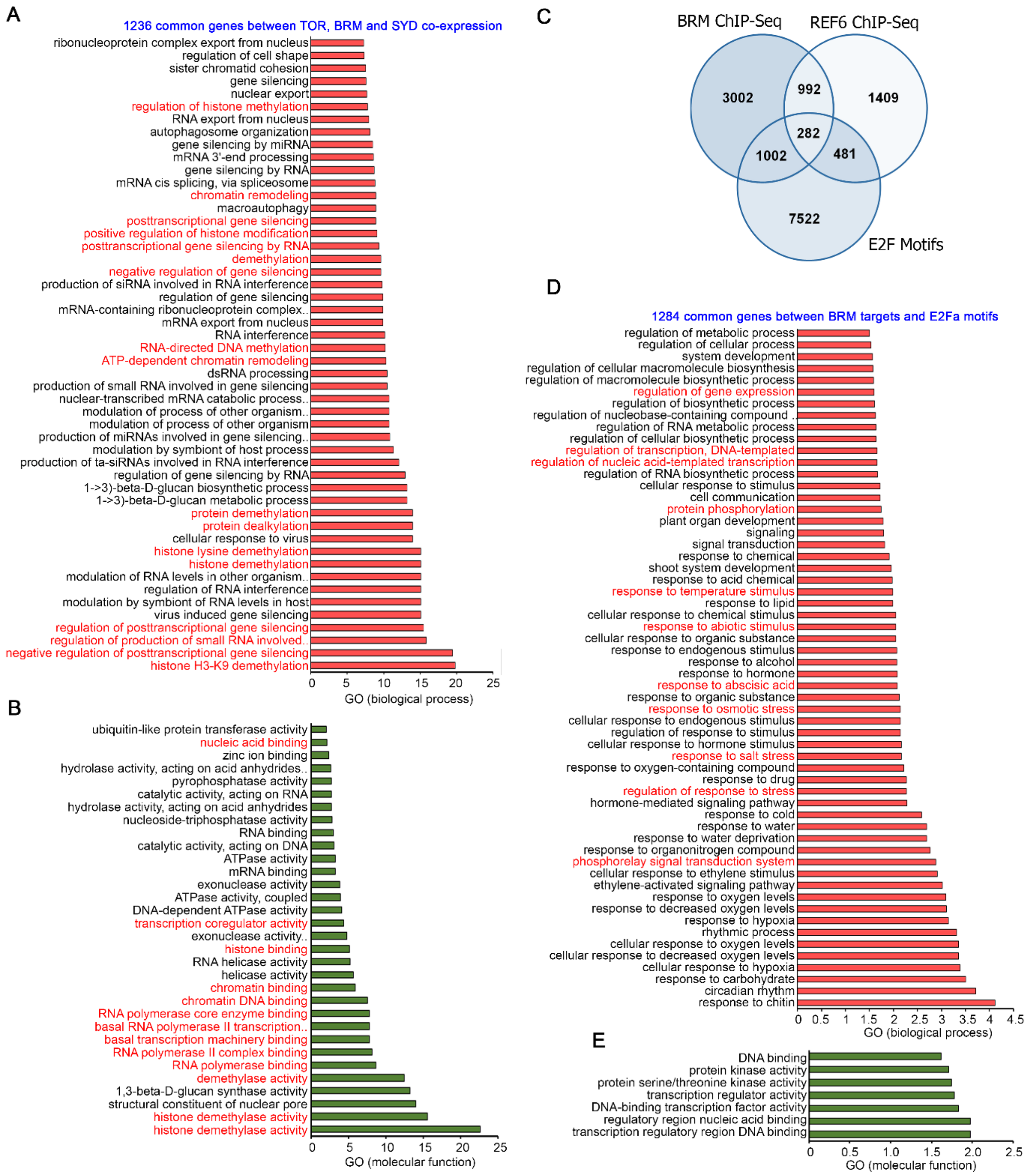


**Supplemental Figure S19. Arabidopsis TOR co-expression shares huge overlap with chromatin remodeler BRAHMA and SYD (CHR3). A,** GO biological process enrichment of 1236 genes commonly co-expressed between TOR, BRM and SYD. **B,** GO molecular function enrichment of 1236 genes commonly co-expressed between TOR, BRM and SYD. **C,** Venn diagram showing overlap of BRM and REF6 ChIP-seq targets with genes containing E2F binding motifs in their promoters. **D,** GO biological process enrichment of 1284 genes between BRM targets and E2F motifs containing genes. **E,** GO molecular function enrichment of 1284 genes between BRM targets and E2F motifs containing genes. BRM and REF6 ChIP-seq data was obtained from publicly available resources (Li *et al.* 2016). Panther 15.0 tool was used to analyse GO fold enrichment using Bonferroni correction and Fisher's exact test type.
